# ACRC/GCNA is an essential protease that repairs DNA-protein crosslinks during vertebrate development

**DOI:** 10.1101/2023.03.07.531502

**Authors:** Cecile Otten, Marin Kutnjak, Christine Supina-Pavic, Ivan Anticevic, Vanna Medved, Marta Popovic

## Abstract

DNA-protein crosslinks (DPCs) are toxic DNA lesions that block all DNA transactions including replication and transcription, and the consequences of impaired DNA-Protein Crosslink Repair (DPCR) are severe. At the cellular level, impaired DPCR leads to the formation of double strand breaks, genomic instability and cell death, while at the organismal level, it is associated with cancer, aging and neurodegeneration. Despite its importance, the mechanisms of DPCR at the organismal level are largely unknown. Proteases play a central role in DPCR, as they remove proteinaceous part of the DPCs, while the peptide remnant crosslinked to DNA is subsequently removed by other repair factors. We characterized the role of putative protease ACRC/GCNA (ACidic Repeat Containing/Germ Cell Nuclear Antigen) in DPCR at the organismal level. For this purpose, we have created new animal models with CRISPR/Cas system: two zebrafish lines with inactive Acrc. We were able to overcome the early embryonic lethality caused by Acrc inactivation by injecting Acrc-WT mRNA and have created a viable animal model to study the role of Acrc in adult tissues. We identified histone H3, topoisomerases 1 and 2, Dnmt1, Parp1, Polr3a and Mcm2 as DPC substrates of Acrc. We have proven that Acrc is absolutely essential for vertebrate development, and that the mechanism behind it is DPC proteolysis.

## Introduction

DNA-protein crosslinks (DPCs) are the second most common DNA lesions in cells (Tretyakova et al. 2015). They occur endogenously at high frequency (6000 DPCs per cell daily) and are induced by byproducts of normal cellular processes such as reactive oxygen species, formaldehyde, and helical DNA changes (Vaz et al. 2017; Ruggiano and Ramadan 2021). External sources of DPCs include UV and IR radiation, as well as various chemicals present in the environment, including chemotherapeutics and transition metals (Tretyakova et al. 2015). Due to their bulkiness, DPCs can stall replication and transcription machineries, ultimately leading to accumulation of mutations and/or cell death. Defective repair of DPCs can lead to aberrations during replication, resulting in genome instability, which in turn can lead to cancer or neurodegeneration (Fielden et al. 2018). DPCs are highly heterogenous and vary by the size of the protein or protein complex, the chemical nature of the crosslink and the DNA topology in the vicinity of the crosslink (intact DNA, double or single strand break). The main DPC repair pathways are (1) proteolysis followed by removal of the peptide remnant from the DNA backbone by the action of nucleotide excision repair (NER) or the tyrosyl-DNA phosphodiesterases TDP1 and TDP2 and (2) the nucleolytic pathway, by which the part of the DNA to which the crosslinked protein is bound is excised (Ruggiano and Ramadan 2021; Vaz et al. 2017; Zhang et al. 2020).

The first DPCR protease, Wss1 (Weak Suppressor of Smt3) was identified in yeast in 2014 (Stingele et al. 2014) followed by SPRTN (SprT-Like N-Terminal Domain), its functional homolog in metazoans (Vaz et al. 2016; Stingele et al. 2016; Lopez-Mosqueda et al. 2016; Maskey et al. 2017). Both proteases belong to zinzin family of metallopeptidases and share a highly conserved protease core (HEXXH) with a catalytic glutamate and three zinc-binding histidines, within the SprT-like domain. SPRTN is an essential protease as its knock-out in mouse causes early embryonic lethality (Maskey et al. 2017); it is specifically active during replication and is very pleiotropic, i.e. can proteolyse various DPCs including histones, topoisomerases 1 and 2 (TOP1 and TOP2), high mobility group protein 1 (HMG1), helicase like transcription factor (HLTF) and Fanconi-associated nuclease 1 (Fan1) (Vaz et al. 2016; Stingele et al. 2016; Lopez-Mosqueda et al. 2016; Mórocz et al. 2017; Weickert et al. 2023). Other proteases which have been recently indirectly implicated in DPCR, but lacking a SprT-like domain, are FAM111A, which interacts with PCNA and plays a role in the resolution of Topo-1cc and trapped PARP-1 (Kojima et al. 2020; Hoffmann et al. 2020), and DNA Damage Inducible 1 (DDI1/Dd1) and 2 (DDI2/Ddi2) which interact with the proteasome and promote replication fork restart after replication stress (Dirac-Svejstrup et al. 2020; Serbyn et al. 2020; Yip et al. 2020; Nowicka et al. 2015; Kottemann et al. 2018; Svoboda et al. 2019). These proteases, like SPRTN, are also specifically active during replication. Interestingly, the acid repeat-containing protein also known as germ cell nuclear antigen (ACRC/GCNA) also harbors a SprT-like domain carrying HEXXH active site (Fielden et al. 2018; Carmell et al. 2016) and has been linked to the repair of topoisomerase 2 (TOP2)-DPCs and DNA-methyltransferase 1 (DNMT1)-DPCs (Dokshin et al. 2020; Bhargava et al. 2020; Borgermann et al. 2021). However, direct evidence of the ACRC protease activity is still lacking, as well as the molecular characterization of its role in DPCR. Given the paucity of knowledge about DPCR on the organismal level and the role of ACRC in DPCR, we turned to zebrafish (*Danio rerio*), a well-characterized vertebrate model organism, to determine the function of Acrc *in vivo*, as well as to biochemically characterize its domains and role in DPCR. Previous studies have shown that ACRC/GCNA plays a role in germ cells: Gcna-1 dysfunction in *C. elegans* resulted in germline lethality (Borgermann et al. 2019a) while *gcna* mutants in Drosophila and zebrafish showed maternal-effect lethality due to chromosome segregation defects in oocytes (Dokshin et al. 2020; Bhargava et al. 2020).

ACRC is an evolutionarily ancient protein that typically harbors four domains: an intrinsically-disordered region (IDR), a Zinc finger and an HMG box for DNA binding, and the SprT-like domain, a Zinc-metalloprotease domain with a characteristic HExxH catalytic core (Carmell et al. 2016; Vaz et al. 2017). Interestingly, the mouse ortholog of ACRC lacks the SprT-like domain and consists almost entirely of the IDR. Male *Acrc* knock-out mice are sterile, suggesting an important role for the IDR domain in mouse male fertility (Carmell et al. 2016; Bhargava et al. 2020; Ribeiro and Crossan 2022). In humans, defective DPCR proteases have been associated with cancer: mutations in *SPRTN* cause a Ruijs-Aalfs syndrome (RJALS) characterized by progeria and early-onset liver cancer due to genomic instability resulting from DPC accumulation (Lessel et al. 2014; Vaz et al. 2016; Halder et al. 2019). Mutations in *ACRC* have been associated with pediatric germ cell tumors (Bhargava et al. 2020) and spermatogenic failure and male infertility (Hardy et al. 2021).

In order to investigate the role of ACRC in DPC repair we have created a separation-of-function mutant zebrafish strain with a mutation in the protease core of Acrc. We show that this catalytic mutation causes early onset embryonic lethality, thus revealing that the proteolytic function of ACRC is essential in early vertebrate development. We confirmed that this effect is specific with a series of rescue experiments where we inject Acrc-WT mRNA into the mutant zygotes which resulted in a complete rescue, i.e. viable embryos, while the injection of the ACRC-E451 mRNA (catalytic mutation) could not overturn lethality phenotype. Mouse ACRC, which lacks a SprT-like domain, could also not rescue the lethality phenotype and neither could SPRTN protease. We then characterized different domains of Acrc, including IDR region, N-terminal part of IDR and C-terminal and found that all three are dispensible for the function of Acrc in the early vertebrate development, while the functional protease core is essential. Using biochemical analyses of the crosslinked proteins we further show DPCs hugely accumulate before the onset of lethality, suggesting that DPC overload due to the inactive ACRC protease is the cause of early embryonic lethality. We detected a significant increase in high molecular weight DPCs, followed by crosslinked proteins of medium and low molecular weight. Specifically, very high accumulation of histone H3-, Dnmt1-, toposiomerases 1 and 2 (Top1-, Top2-), Parp1-, Polr3a- and Mcm2-DPCs in Acrc mutant fish strongly suggest that latter proteins are newly identified DPC substrates of Acrc protease. Furthermore, by injecting Acrc-WT mRNA in homozygous mutant strain we have been able to compensate for the need of Acrc in the early vertebrate development, and the adult fish are viable. Thus, we have created a viable animal model to study the role of Acrc in adult organism. We show that human and zebrafish Acrc share 1-to-1 orthology, conserved synteny and domain organization. We also shown that acrc is highly expressed across adult tissues, with predominant expression in gonads. The consequences of the impaired Acrc activity in the adult tissues remain to be determined. Our results demonstrate the importance of Acrc for DPC repair at the organismal level and the importance of the protease core for Acrc function in early vertebrate development. Furthermore, we demonstrate that besides SPRTN protease, there is another essential metalloprotease which can resolve a wide range of DPC substrates and has potential therapeutic implications, especially given the fact that it is involved in the repair of topoisomerase- and Parp1-mediated DNA damage.

## Results

### ACRC/GCNA is a highly conserved protein with a SprT-like domain

The ACRC protein is highly conserved with one-to-one orthology throughout the animal kingdom (Figures 1 and S1-3). Except for a subset of rodents that includes the house mouse (*Mus musculus*) and the rat (*Rattus norvegicus*), all vertebrate and invertebrate orthologs have a SprT-like domain (Figures 1 and S2). As the SprT-like domain is a likely active protease domain which could have a function in DPCR, we specifically analyzed the evolutionary conservation and protein structure of this domain. The 3D Alphafold structural model shows a high degree of conservation between the human and zebrafish SprT-like domains. Both domains were modelled with very high or high degree of confidence (per-residue confidence score, pLDDT > 90 and 70-90, respectively) (Figure 1A). The ACRC protease core comprises three α-helices and three β-sheets carrying the active site, which consists of a catalytic glutamate (E593 in human and E451 in zebrafish) and three Zn^2+^-binding histidines (H592, H596 and H609 in human and H450, H454 and H467 in zebrafish) (Figure 1A). Three β-sheets are conserved at the C-terminal end of the SprT-like domain in both species. Both proteins also share a long unstructured and intrinsically disordered region (IDR) in the N-terminal (Figures 1A and S5B). Interestingly, the human and zebrafish orthologs differ strongly in their protein charge clusters: human ACRC carries a large negative (Asp/Ser-rich) and a large positive charge cluster (Lys/Arg-rich) within the IDR region, whereas zebrafish Acrc has no significantly charged regions (Figure 1A). In addition, the nature of the amino acid repeats within the IDR differs: human ACRC contains many periodic repeats spanning a large portion of the IDR, whereas zebrafish Acrc has only two short repeats (R1 and R2) (Figure 1A). The Alphafold model of the C-terminal part of zebrafish Acrc predicts two α- helices with high to very high confidence, as well as intermittent structured regions (low to high confidence) within the otherwise largely disordered N-terminal part. In comparison, the human ACRC has a shorter, unstructured C-terminus and is largely disordered in the N-terminal part (Figures 1A and S5B). Interestingly, when we compared the human and zebrafish protein regions outside the SprT-like domain with their mouse ortholog, we observed similar large unstructured and disordered regions, including periodic repeats, whereas the charge clusters of mouse ACRC are more similar to its human ortholog (Figures 1 and S5B).

**Figure 1.**
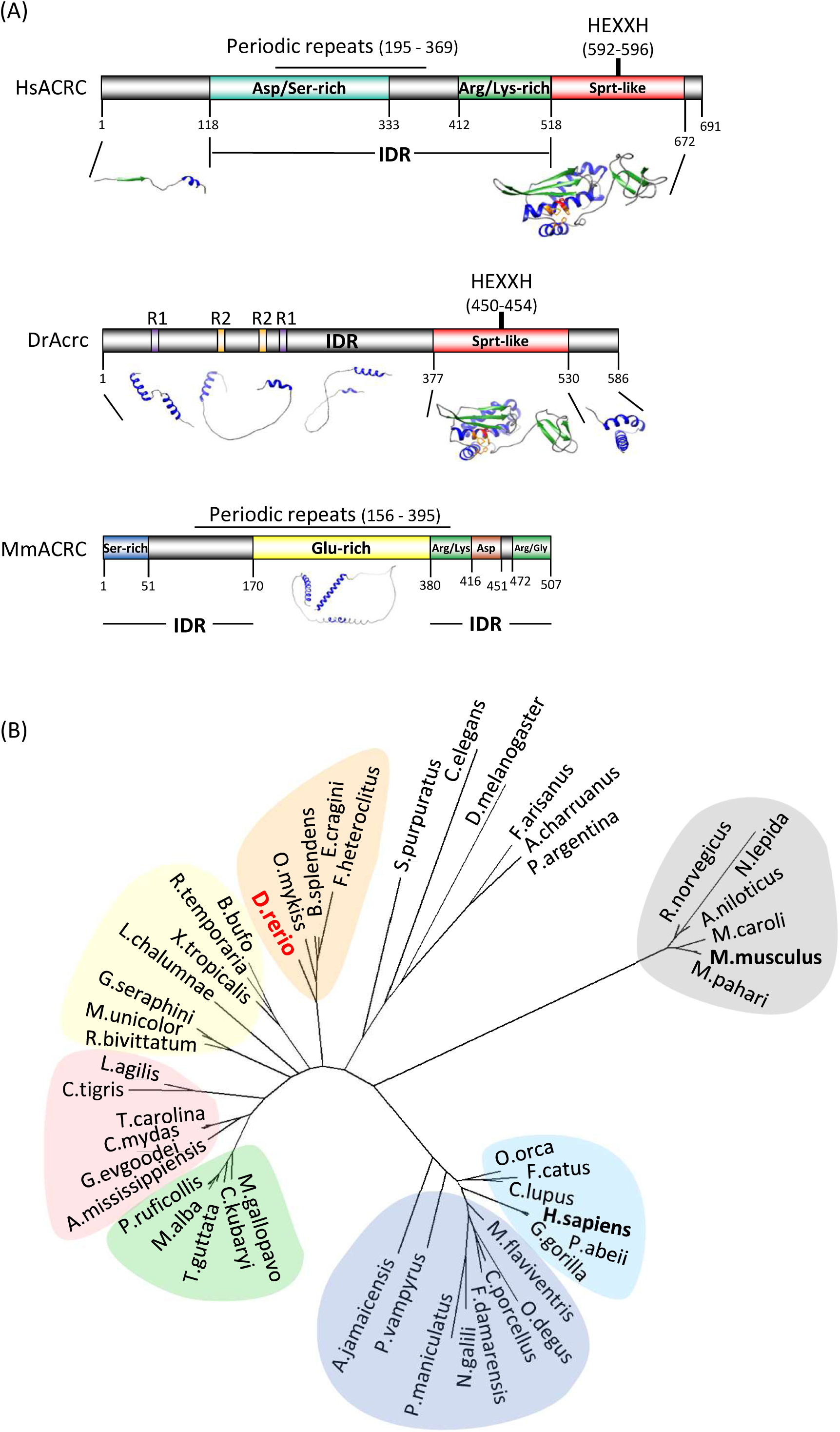
ACRC is an evolutionary conserved putative protease. **(A)** Comparison of human, zebrafish and mouse ACRC/GCNA protein domains and motifs and structural model of human ACRC. Domains and motifs were identified using SAPS and Motif Scan workspace. Models were build using Alphafold and analyzed with Chimera. HEXXH, protease core; IDR, Intrinsically Disordered Region; R, repeats. The metalloprotease core of ACRC includes the active glutamate shown in red and Zn^2+^ binding histidines shown in orange. **(B)** Phylogenetic analysis of ACRC/GCNA orthologs. Full-length (LG 4) protein sequences were aligned using MAFFT and the phylogenetic tree was constructed in PhyML. Blue: non-rodents mammals, dark blue: rodents and bats with a SprT-like domain, grey: rodents without SprT-like domain, green: birds, red: reptiles, yellow: amphibians, orange: fish, colorless: invertebrates.

When we performed a phylogenetic analysis using the full-length protein sequences, the rodent subgroup lacking the SprT-like domain clustered separately from other mammalian orthologs including humans and other rodents (Figure 1B). As expected, when we used the protein alignment of ACRC SprT-like domains, the phylogenetic tree was very similar to the analysis when the complete protein sequences were used (Figures S1 and S2). However, when we performed a phylogenetic analysis using the N-terminal domains and excluding SprT-like domains, all mammalian orthologs clustered together, indicating that the rodent subgroup is evolutionarily close to other mammalian groups with a SprT-like domain (Figure S3, Table S1).

Human and mouse *ACRC* genes are located on the X chromosome, whereas zebrafish *acrc* is positioned on chromosome 14 (Figure S4). Of note, zebrafish do not have sex chromosomes, but one of the chromosome clusters that determine gender is located on chromosome 14 (Howe et al. 2013). Otherwise, the gene environments of the human and mouse *ACRC* orthologs are quite similar and partially conserved with the gene environment of zebrafish *acrc*.

In summary, phylogenetic, syntenic and domain analyses showed that zebrafish is a good model to study ACRC function because it shares many features with human ACRC, including the SprT-like domain that the mouse ortholog lacks.

### *Acrc* and *Sprtn* are highly expressed during embryonic development and in adult tissues

We determined the mRNA expression of *acrc* in adults and embryos and compared it with the expression of *sprtn* to gain more insight into these two DPC proteases. To facilitate comparison of expression levels, we set arbitrary thresholds following previous publications (Lončar et al. 2016) (Anticevic et al. 2024, 2023): high expression when MNE is greater than 60 × 10^6^ (Ct values < 22), moderate when MNE is 2 × 10^6^ – 60 × 10^6^ (Ct = 23–26) and low when MNE is < 2 × 10^6^ (Ct > 27). *Acrc* shows very high expression in the gonads, particularly in the ovaries, where it is expressed 5.6-fold more than *sprtn* (Figures 2A and 2B). In comparison, the expression of *sprtn* in the testes is very high: 9-fold higher than that of *acrc*. In all other tissues examined, including brain, liver, kidney, intestine and gill, we found moderate to high expression of both genes (Figures 2A and 2B).

**Figure 2.**
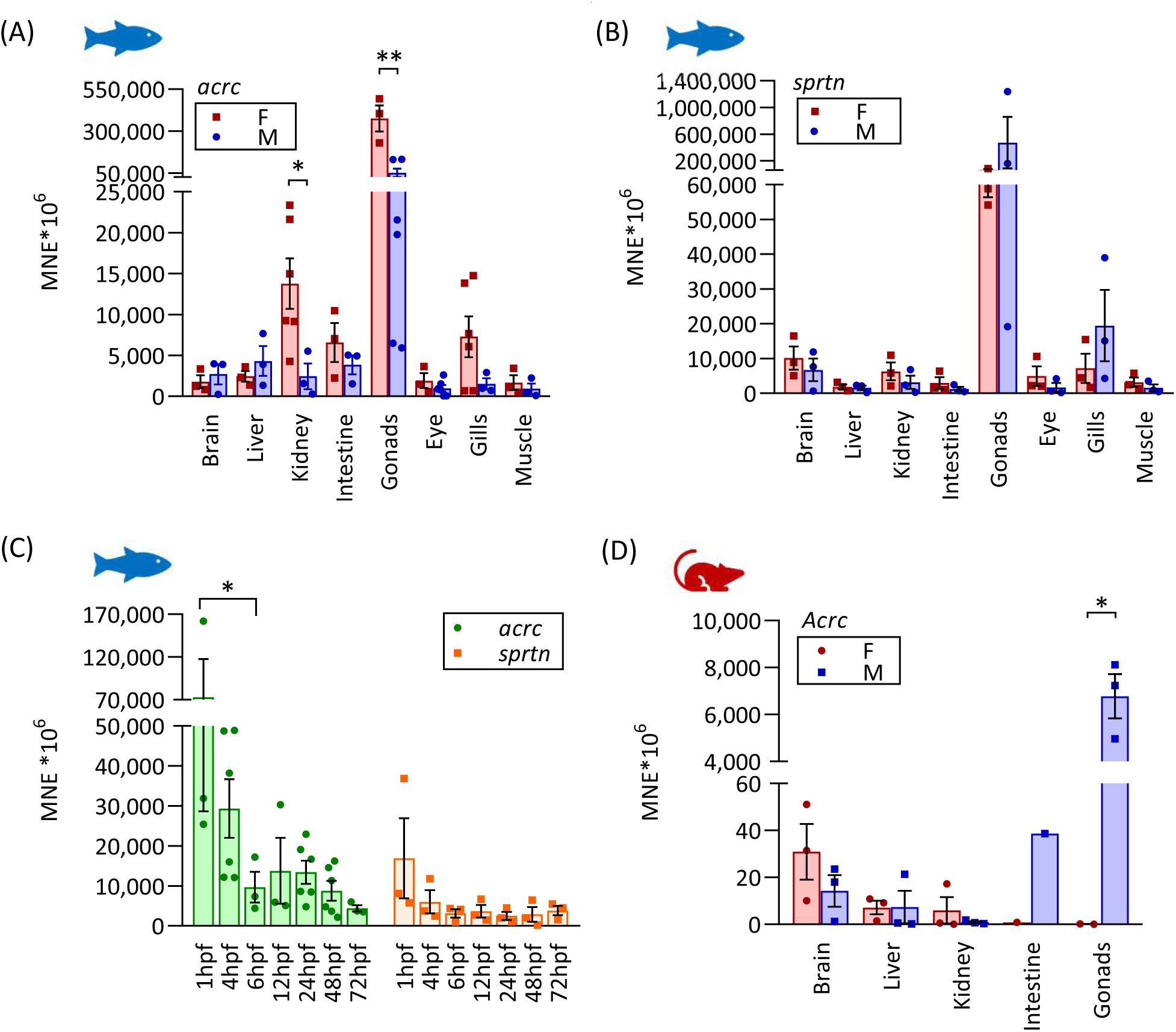
ACRC and SPRTN are highly expressed during embryonic development and in adult tissues. **(A)** Expression levels of *acrc* and **(B)** *sprtn* in tissues of adult female (F) and male (M) zebrafish; **(C)** Comparison of *acrc* and *sprtn* expression levels in zebrafish embryos during development (hpf, hours post fertilization); **(D)** Expression levels of *Acrc* in tissues of adult mice quantified by qRT-PCR. Data represent MNE (Mean Normalized Expression) ± SD normalized to the housekeeping genes *elongation factor 1α (eef1a1l1*) for zebrafish and *Rplp0* for mouse.

Next, we determined the expression levels of *acrc* and *sprtn* during the embryonic development when replication and transcription rates are very high and accumulated DPCs can be especially detrimental. Gene expression levels of *acrc* and *sprtn* are very high in zebrafish embryos between 1 and 72 hours post-fertilization (hpf) (Figure 2C). Both genes showed the highest expression at 1 hpf, indicating maternal mRNA deposition in the egg, as zygotic transcription is only fully activated at 3 hpf and maternal transcript levels decrease between 2 and 3 hpf (Tadros and Lipshitz 2009; Laue et al. 2019). Expression levels of *acrc* decline rapidly until 6 hpf and remain approximately stable until 2 days post-fertilization (dpf) before decreasing further. In comparison, expression levels of *sprtn* also decrease rapidly until 6 hpf, but remain stable until 3 dpf. *Acrc* levels were at least three times higher than *sprtn* levels at all stages until 2 dpf, indicatinga crucial role during development. It is important to note that the expression levels of both genes are still very high at 72 hpf (MNE 5,000*10^6^ ) (Figure 2C).

To further investigate the role of the SprT-like-domain, we compared the gene expression levels of zebrafish *acrc* (which has a SprT-domain) and mouse *Acrc* (which lacks the SprT domain) in adult tissues. The expression pattern differs markedly between the two genes, with zebrafish *acrc* being moderately to highly expressed in all examined tissues, with the highest expression in ovary and testis, whereas mouse *Acrc* is dominantly expressed in testis, and at low to moderate levels in other tissues (Figures 2A and D).

### Acrc mutations in the SprT-like domain cause early embryonic lethality in zebrafish

To study DPC repair *in vivo*, we generated two zebrafish *acrc* mutant strains targeting the SprT-like domain using the CRISPR/Cas9 system (Figures 3A and S5A). One mutant strain carries an in-frame deletion of 12 nucleotides in the putative catalytic site resulting in the deletion of the catalytic glutamate E451 and of the following three amino acids in the protease core, including the Zn bearing Histidine H454 (ΔEMCH). We named this mutant allele *rbi5* (*ruder boskovic institute* 5) following naming nomenclature suggested by ZFIN (The Zebrafish Information Network, zfin.org). The second mutant strain carries mutations in exon 12, leading to frameshift and premature stop codons at the 477. and 472. amino acid positions (Figure S5A). We named these mutant alleles *rbi8* and *rbi9* (*ruder boskovic institute* 8 and 9). The phenotype of both types of mutant strains is maternal zygotic embryonic lethal (Figures 3C and S6C). Most importantly, the fact that the catalytic mutation (rbi5, ΔEMCH) in the protease core causes embryonic lethality (Figure 3) strongly suggests that it is the proteolytic function of Acrc that is responsible for the observed phenotype. Furthermore, embryos with homozygous mutant mothers show an early lethality phenotype, whereas homozygous mutant embryos with heterozygous mothers show no morphological defects (Figures 3D-F) which demonstrates the requirement for maternal deposition of functional Acrc protein and/or mRNA in the eggs.

**Figure 3.**
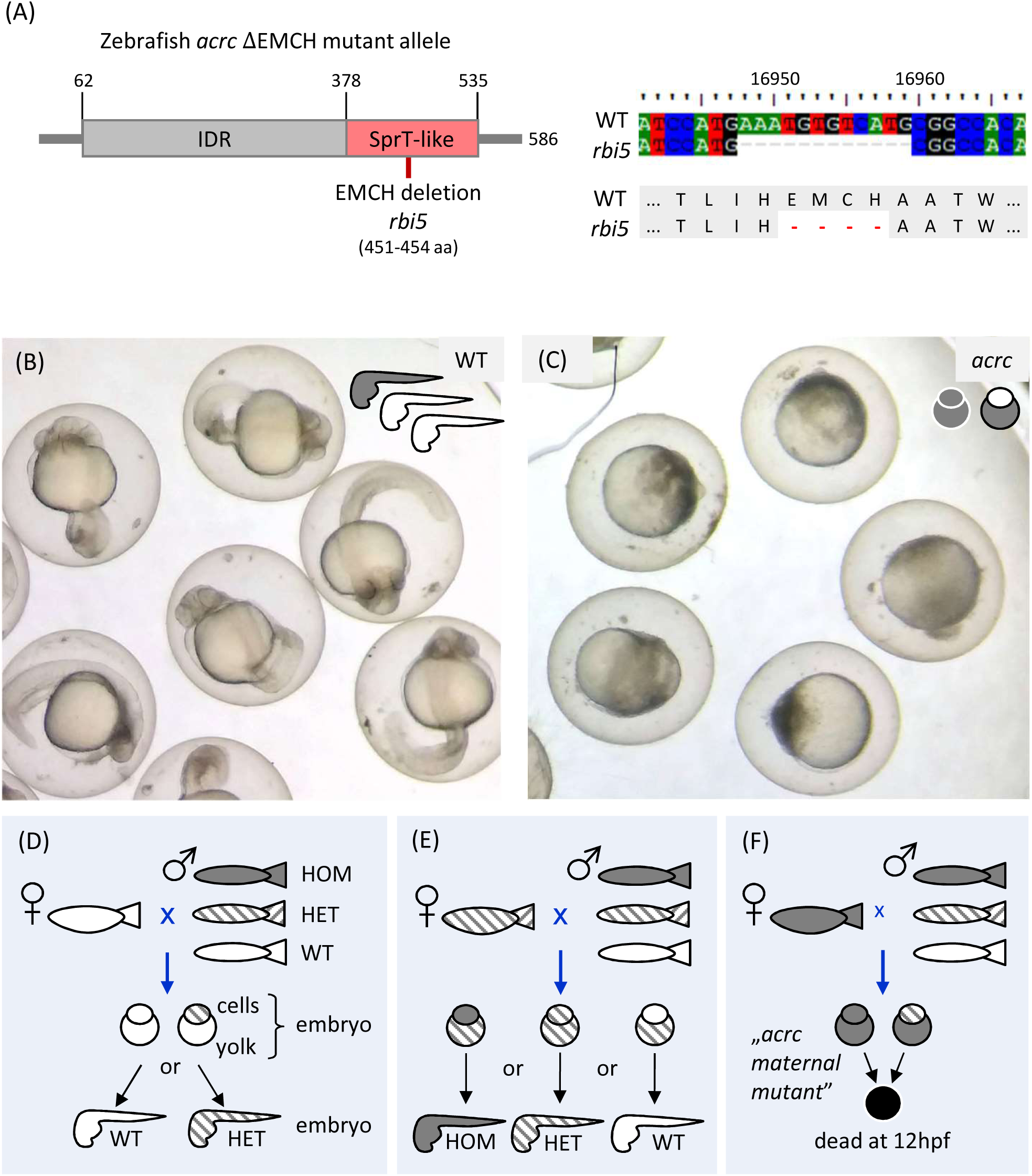
Creation of *acrc* mutant zebrafish strain carrying a catalytic mutation. **(A)** Scheme of the Acrc protein and the position of the 4 amino-acids deletion (ΔEMCH, 451-454 aa) in the enzymatic core of the SprT-like domain in the *rbi5* allele, which was created using CrispR/Cas9 technology; (B) Representative pictures of zebrafish embryos - WT and (C) *acrc ^rbi5/rbi5^* maternal mutants (ΔEMCH) at 24 hpf (hpf, hours post fertilization); (D - F) Schemes representing the genotype-phenotype correlations upon loss of one or both *acrc* wt alleles: D – crossing a WT female with any male (WT: white, heterozygote: stripes, or mutant: grey) always leads to viable fish, E - crossing a heterozygous female with any male (WT, heterozygote or mutant) always leads to viable fish, and F - crossing an *acrc* mutant female with any male (WT, heterozygote or mutant) always results in embryos with an early lethal phenotype (black yolk).

### The putative catalytic core of the SprT-like domain is essential for Acrc function during early embryonic development

To confirm that the catalytic mutation is responsible for the lethal phenotype and to gain insight into the function of other Acrc domains, we injected *in vitro* transcribed and capped mRNA encoding the wild-type or mutated versions of Acrc into *acrc^rbi5/rbi5^* maternal mutant embryos at the 1-cell stage and examined whether they could compensate for the loss of Acrc and thus “rescue” the early lethality phenotype of the *acrc* mutant. Each *in vitro* synthesized mRNA was injected into WT and *acrc* maternal mutant embryos, and the resulting phenotypes were scored according to severity in 5 categories of overall abnormality (Figures 4 and S6). This was performed in both *acrc^rbi5/rbi5^*maternal mutant embryos (Figure 4, Table S2) and in *acrc^rbi8/rbi9^* maternal mutants (Figure S6, Table S3).

**Figure 4.**
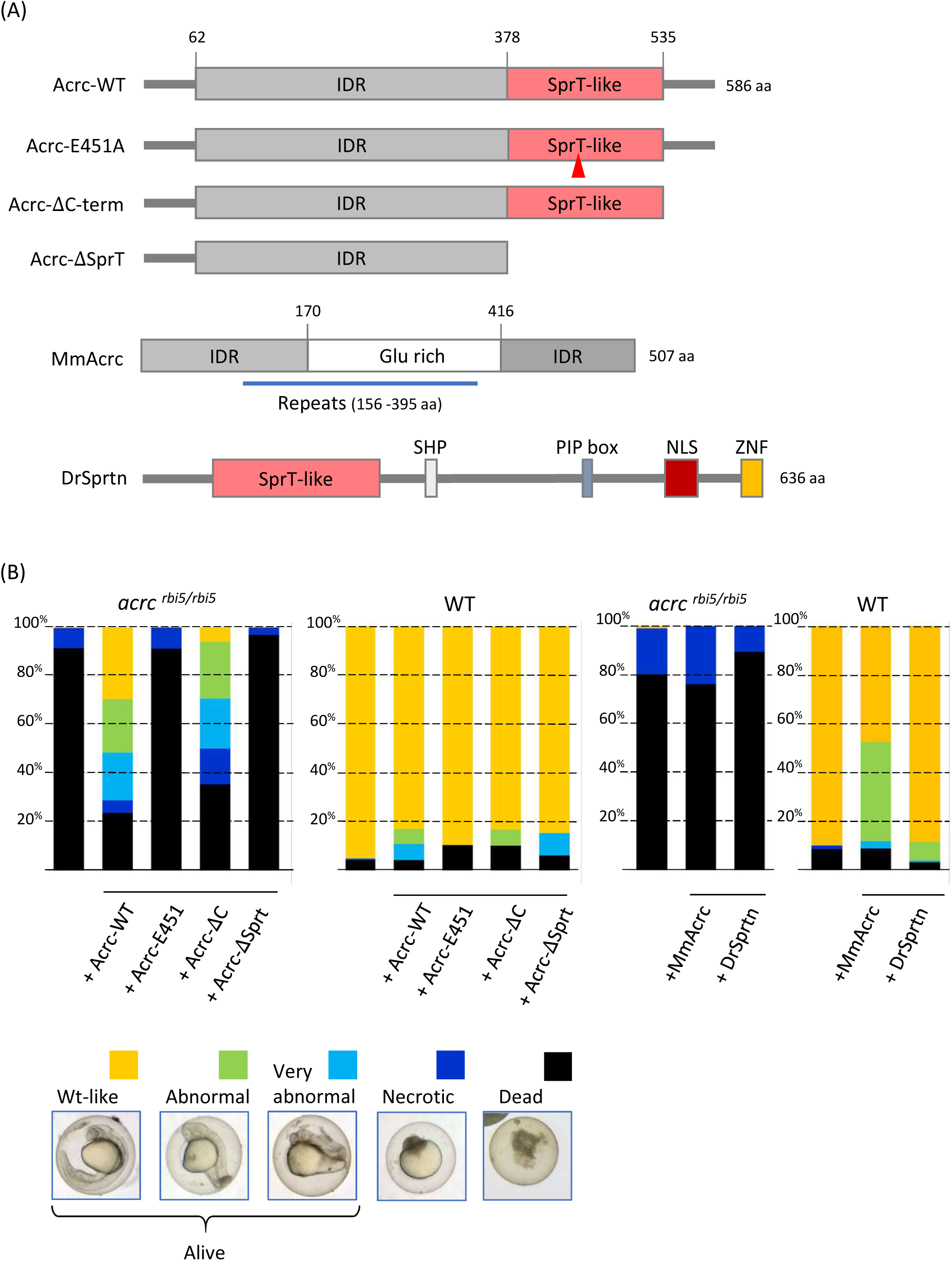
Injection of ACRC-WT mRNA fully rescues the *acrc* maternal mutant phenotype, unlike the injection of ACRC-E451A, DrSprtn and MmAcrc mRNA. **(A)** Scheme of the injected rescue constructs: Acrc-WT, Acrc-E451A, Acrc-ΔC-term, and Acrc-ΔSprT; Mm, mouse musculus; Dr, Danio rerio **(B)** Phenotype characterization of rescued embryos with representative pictures. WT and *acrc^rbi5/rbi5^* maternal mutant embryos were injected with 1 nl of 250 ng/ml mRNA derived from the rescue constructs. Phenotypes were scored at 1 dpf and divided into 5 categories: #1: dead, #2: necrotic, #3: very abnormal, #4: abnormal, and #5: WT-like. Embryos in categories #3, 4 and 5 were considered alive. Data shown are means. For detailed results, see Supplemental Tables 2-4.

Firstly, we injected an Acrc-WT mRNA into mutant embryos and observed a rescue of the lethal phenotype (Figures 4 and S6). 71,2% of *acrc^rbi5/rbi5^* maternal mutant embryos injected with Acrc- WT mRNA were alive after 24 hpf (n = 121 in 4 independent experiments), while 94,8% of *acrc^rbi8/rbi9^* maternal mutants were alive after injection with Acrc-WT mRNA (n = 475 in 5 independent experiments) (Tables S2-4). The degree of rescue was such that in fact, the majority of injected embryos (70-90%) develop normally to adulthood (3 months old) and older age (1.5 year old). An example of uninjected and injected *acrc^rbi8/rbi9^* maternal mutants at 1dpf can be seen in Figure S6C.

Secondly, we confirmed that the catalytic mutation in the putative protease core is essential for embryonic development, as the injected Acrc-E451A mRNA could not rescue the lethal phenotype (Figures 4 and S6). None of the *acrc^rbi5/rbi5^* maternal mutant embryos injected with Acrc-E451A mRNA was alive after 24 hpf (n = 28 in 2 independent experiments). The same was observed after the injection of Acrc-E451A mRNA in the *acrc^rbi8/rbi9^* maternal mutants (n=193 in 4 independent experiments), suggesting that the glutamate 451 located in the protease core is crucial for Acrc function during early embryonic development (Figures 4B and S6).

Next, we tested whether the other domains of Acrc affect the lethality phenotype. To this end we tested the function of a truncated version of Acrc lacking the C-terminal tail (Acrc-ΔC-term) and of Acrc lacking SprT domain (Acrc-ΔSprT). The construct with small C-terminal truncation, which has intact SprT and IDR domains (Acrc-ΔC-term), was able to rescue the lethal phenotype, similar to Acrc-WT (Figures 4 and S6, Tables S2-4). 50% of *acrc^rbi5/rbi5^* maternal mutant embryos injected with Acrc-ΔC-term mRNA were alive after 24 hpf (n = 17, 1 experiment), while 92,5 % *acrc^rbi8/rbi9^* maternal mutants were alive after injection with the Acrc-ΔC-term mRNA (n = 119 in 3 independent experiments) (Tables S2-4). The Acrc construct lacking the SprT domain (Acrc-ΔSprT) was not able to rescue the lethal phenotype (n = 31 in 1 experiment for *acrc^rbi5/rbi5^*, n = 109 in 2 experiments for *acrc^rbi8/rbi9^* maternal mutants) (Figures 4 and S6). The stability of the injected mRNA constructs was verified by RT-PCR (Figures S6D-E).

Overall, the rescue experiments clearly demonstrate that Acrc is an active protease whose protease activity is absolutely required for survival at early embryonic stages. Remarkably, *acrc* maternal mutant embryos of both strains (*rbi5* and *rbi8/9*) injected with full-length Acrc-WT mRNA reached adulthood and were fertile, allowing us to breed and maintain the *acrc* mutant lines.

### Mouse Acrc did not compensate for loss of zebrafish *acrc* during early embryonic development

To further confirm that the proteolytic function of ACRC is essential for the early development, we tested whether mouse ACRC, which lacks a SprT-like domain (Figure 1A), can rescue early embryonic lethality in *acrc* mutant fish (*acrc^rbi5/rbi5^*). To this end, we injected mouse *ACRC* mRNA into zebrafish *acrc^rbi5/rbi5^* mutant embryos (Figure 4). None of the injected embryos were alive at 24 hpf (n = 105 in 2 independent experiments) (Figure 4B), confirming that the Acrc protease activity is required for its function in early development, rather than the other Acrc domains. Control injections confirmed that injecting mouse *ACRC* mRNA is not toxic for zebrafish embryos (Figure 4B). 91% of WT embryos injected with MmAcrc-WT mRNA were alive at 1 dpf (n = 107 in 2 independent experiments). In addition, RT-PCR confirmed that MmAcrc-WT mRNA remains stable at least for 2 days after the injection (Figure S6E).

### Sprtn did not compensate for loss of zebrafish *acrc* during early embryonic development

Considering that the protease Sprtn, like Acrc, has a SprT-like domain (Figure 4A) and that they have very similar protease cores (Fielden et al. 2018c), we tested whether Sprtn can compensate for the absence of Acrc during the early development. Interestingly, injection of zebrafish Sprtn mRNA into *acrc^rbi5/rbi5^* maternal mutant embryos failed to rescue the early lethality phenotype and not a single viable embryo was present at 1 dpf post injection (n = 105 in 2 independent experiments) (Figure 4B, Table S4), highlighting the unique function of Acrc during early embryonic development. The stability of the injected mRNA constructs was verified by RT-PCR (Figure S6D).

### *Acrc* mutants accumulate cellular DPCs before the onset of lethality

After we confirmed that the proteolytic function of Acrc is essential during early embryonic development, we investigated whether the molecular mechanism behind the embryonic lethality is impaired DNA-protein crosslink repair.

To this end, we isolated DPCs from *acrc^rbi8/9^* mutant embryos at 6 hpf when embryos still exhibited normal WT-like morphology, before the onset of necrosis. Using the SDS-KCl method, we precipitated protein-bound DNA to separate it from protein-free DNA. Thus, the amount of DNA in the SDS pellet is used as a measure of DNA-protein crosslink amount expressed as a percentage of crosslinked DNA (Zhitkovich and costa 1992; Mórocz et al. 2017). *Acrc^rbi8/^*^9^ mutants (ACRC-ΔC) accumulated significantly more DPCs in comparison to WT embryos at 6 hpf: the percentage of crosslinked DNA relative to free DNA in *acrc^rbi8/9^* mutants was 32.75 ± 2.88%, in comparison to 18.04 ± 3.09% in WT embryos (Figure 5A).

**Figure 5.**
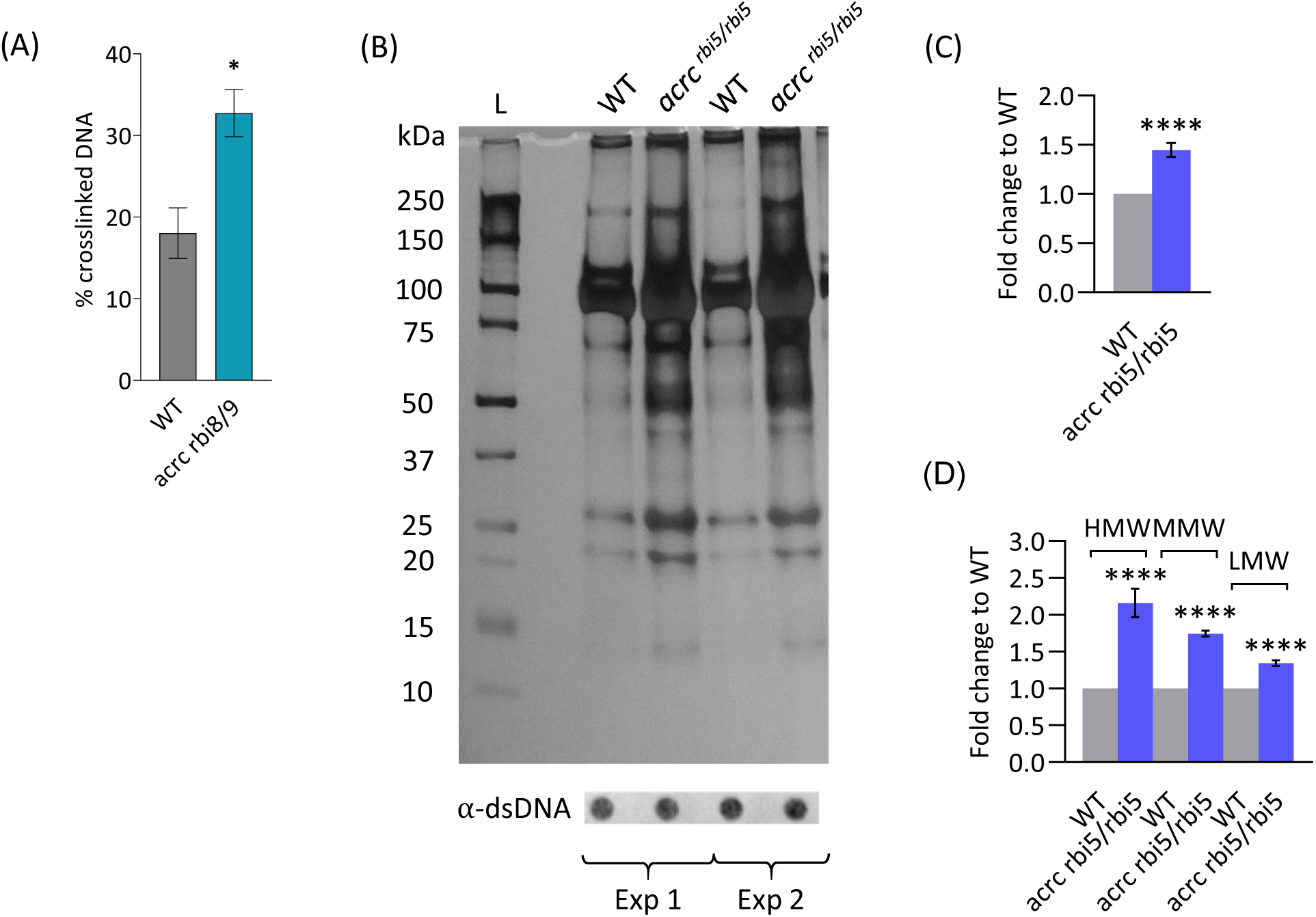
DPC levels are significantly increased in *acrc* mutant embryos at 6 hpf. **(A)** Total DPC levels (% of crosslinked DNA) isolated using the KCl/SDS method in 6 hpf WT and *acrc^rbi8/9^* mutant embryos. An unpaired Student’s t-test was performed for 3 biological replicates, pools of 100 embryos each (*: p-value = 0.0254); **(B)** Total DPC levels (crosslinked proteins) isolated using modified RADAR method from 6 hpf WT and *acrc^rbi5/rbi5^* embryos in two biological replicates. Total DPCs were isolated from n = 50 embryos each for WT and *acrc^rbi5/rbi5^*, resolved on an SDS acrylamide gel, and stained with silver. Dot-blots showing DNA loading controls for DPC analysis prior to benzonase treatment are shown below; a DPC equivalent of 25 ng total DNA was loaded per well; **(C-D)** Quantifications of (B) using ImageJ based on data shown here and in Supplemental Figure S7A (4 biological replicates). An unpaired Student’s t-test was performed for 3 or 4 biological replicates, pools of 50 embryos each (****: p-value < 0.0001). Data shown are means ± SEM. L-ladder; HMW-high molecular weight, MMW-medium molecular weight and LMW-low molecular weight DPCs.

Considering that proteins forming the crosslinks cannot be detected using the KCl/SDS DPC isolation, due to the nature of the protocol (Torrecilla et al. 2024), we performed a more detailed analysis of DPCs using the modified RADAR isolation protocol which we previously adapted for zebrafish embryos (Anticevic et al. 2023). *Acrc^rbi5/rbi5^* mutants (ACRC-ΔEMCH) had significantly increased levels of total cellular DPCs compared with WT embryos (1.44 fold change, n = 4 biological replicates) (Figures 5B-C and S7A). DPCs which accumulated in *acrc^rbi5/rbi5^* mutants consisted of proteins of low (<40 kDa, LMW), medium (41-150 kDa, MMW) and high molecular weight (> 151 kDa, HMW) (Figures 5B, 5D and S7A), indicating that the Acrc protease has multiple DPC substrates.

### The catalytic mutation in the Acrc protease leads to the accumulation of Dnmt1-, Top1-, Top2-, histone H3-, Parp1-, Polr3a- and Mcm2-DPCs

We further used the modified RADAR method for DPC isolation, as it enables the isolation of intact crosslinked proteins and thus subsequent identification of the protein species. We first investigated whether DNA (cytosine-5)-methyltransferase 1 (Dnmt1) and topoisomerase 2 (Top2) are crosslinked in *acrc^rbi5/rbi5^* mutant embryos at 6 hpf when embryos exhibited normal WT-like morphology, as it was previously suggested that these two proteins might be substrates of Acrc (Borgermann et al. 2019; Dokshin et al. 2020). Indeed, in the absence of proteolytically active Acrc, we observed striking accumulation of Top2-DPCs: 9.7 ± 0.32 fold in comparison to WT embryos (n = 2) (Figures 6A-B and S7B). Dnmt1-DPC levels were also significantly higher in the *acrc^rbi5/rbi5^*mutants (2.0 ± 0.36 fold, n = 3) (Figures 6A-B and S7B).

**Figure 6.**
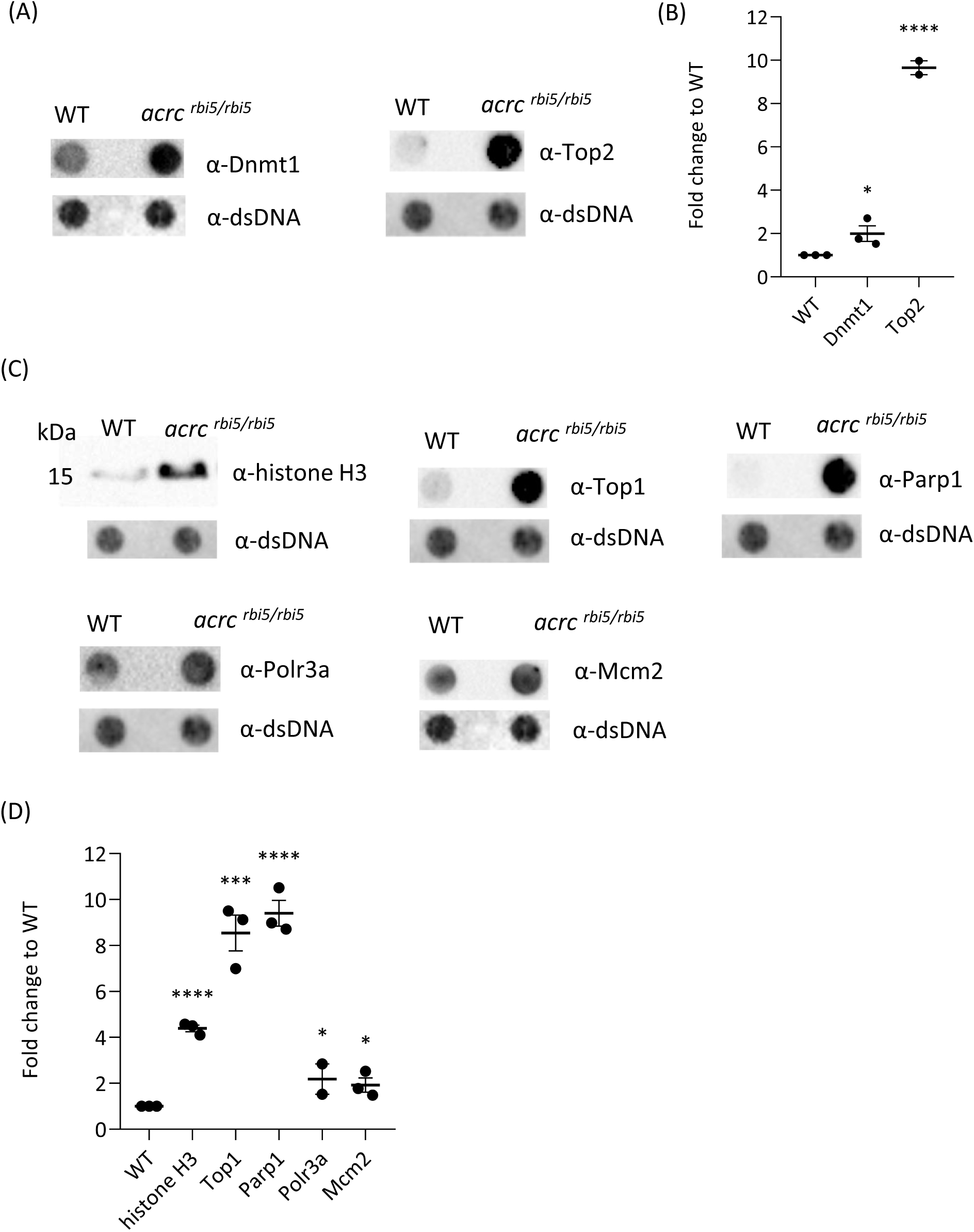
ACRC is involved in the repair of multiple DPC lesions. **(A)** Dnmt1-DPCs and Top2-DPCs accumulate in *acrc^rbi5/rbi5^* mutant embryos. DPCs were isolated by RADAR assays from n = 50 embryos (6 hpf); Dnmt1-DPCs and Top2-DPCs were visualized using dot blot with specific antibodies with a corresponding dot blot for dsDNA as a loading control. A DPC equivalent of 25 ng total DNA was loaded per dot; **(B)** Quantification of (A) using ImageJ based on data shown here and in Supplemental Figure S7B (3 biological replicates); **(C)** Histone H3-DPCs, Top1-, Parp1-, Polr3a- and Mcm2-DPCs accumulate in *acrc^rbi5/rbi5^* mutant embryos. DPCs were isolated by RADAR assays from n = 50 embryos (6 hpf); histone H3-DPCs were detected by western blot, with a corresponding dot blot for dsDNA as a loading control; Top1-, Parp1-, Polr3a- and Mcm2-DPCs were detected with specific antibodies using dot blot (a DPC equivalent of 25 ng total DNA was loaded per dot). Dot blots showing DNA loading controls for DPC analysis prior to benzonase treatment are shown below; 2 ng of total DNA were loaded per well; **(D)** Quantification of (C) using ImageJ based on data shown here and in Supplemental Figure S7C (3 biological replicates). An unpaired Student’s t-test was performed for 2 or 3 biological replicates, pools of 50 embryos each (*: p-value < 0.05, ***: p-value < 0.001, ****: p-value < 0.0001). Data shown are means ± SEM.

Considering that histones are the most abundant DPCs composing approx. 80% of all cellular DPCs (Kiianitsa and Maizels 2020), we tested whether core histones could be DPC substrates of the Acrc protease using histone H3 as a representative. In DPC isolates we specifically detected histone H3-DPC using a H3-specific antibody and found that *acrc^rbi5/rbi5^*mutants with catalytic mutation in the protein core had significantly elevated levels of H3-DPCs (4.4 fold change) in comparison to WT embryos: 4.4 ± 0.15 fold change (n = 3) (Figures 6C-D and S7C). Next, we tested whether Acrc is involved in the removal of other abundant cellular DPCs previously identified by mass spectrometry analysis in human cells (Kiianitsa and Maizels 2020): TOP1 (DNA topoisomerase 1), PARP1 (Poly [ADP-ribose] polymerase 1), POLR3A (DNA-directed RNA polymerase III subunit RPC1) and MCM2 (Minichromosome maintenance complex component 2). Using specific antibodies, we found that *acrc^rbi5/rbi5^* embryos had significantly elevated levels of all of the detected proteins: Top1-DPCs (8.5 ± 0.78 fold change, n = 3), Parp1-DPCs (9.4 ± 0.56 fold change, n = 3), Polr3a-DPCs (2.2 ± 0.66 fold change, n = 2) and Mcm2-DPCs (1.9 ± 0.31 fold change, n = 3) in comparison to WT embryos (Figures 6C-D and S7C).

## Discussion

We report that the Acrc protein is essential during vertebrate embryonic development through its function as a DPC repair protease. A functional SprT-like domain, and particularly a catalytic glutamate, are indispensable for the full functionality of Acrc and thus for embryo survival. Our *in vivo* analysis of different Acrc domains showed that the SprT-like domain is required to rescue the lethality phenotype in *acrc* mutants, while the intrinsically disordered region (IDR) alone is not sufficient. We further showed that Acrc is involved in the repair of multiple cellular DPCs including Dnm1, topoisomerases 1 and 2, histone H3, Parp1, Polr3a and Mcm2.

Given its essential function during vertebrate embryonic development, it is not surprising that ACRC is highly conserved and exhibits one-to-one orthology throughout the animal kingdom (Figures 1 and S1-3). In support of our functional studies, we found that the SprT-like domain in the C-terminal half of ACRC is remarkably conserved from invertebrates to human (Figures 1 and S2), while the N-half of the protein is largely disordered and variable across species (Figure S3). Considering that the mouse ACRC lacks a SprT-like domain (Carmell et al. 2016), we performed an in-depth analysis of rodent ACRC orthologs and showed that many rodent species possess a SprT-like domain; it will be interesting to determine why it has been lost during evolution, specifically in certain mouse and rat species and how this impacted ACRC function in those species (Figures 1B and S1-3, Table S1). We have also shown that the 3D structural models of the SprT-like domain of the human and zebrafish ACRC orthologs are nearly identical (Figure 1A), making zebrafish an excellent model for studying ACRC-mediated DPC proteolysis.

ACRC was originally described as a protein with a specific role in germ cells (Dokshin et al. 2020; Bhargava et al. 2020; Carmell et al. 2016), while another study suggested an important role in somatic cells of *C. elegans* (Borgermann et al. 2019). We therefore investigated mRNA expression of *acrc* in adult zebrafish tissues. We found very high and dominant expression of *acrc* in both ovaries and testes (Figure 2A) consistent with previous work showing that ACRC plays an important role in germ cells (Bhargava et al. 2020; Dokshin et al. 2020 Carmell et al. 2016). Interestingly, we found that *acrc* is also expressed in all other tissues analyzed (Figure 2A). In contrast, we confirmed that mouse *Acrc* is expressed almost exclusively in the testis (see Gene Expression Database for Gcna under www.informatics.jax.org, February, 2023, and Figure 2D), which further highlights the difference between mouse Acrc which lacks a SprT-like domain and other vertebrate orthologs (Figure 1). Of note, the *ACRC* gene expression pattern in humans is more similar to zebrafish *acrc* expression pattern: unlike in mouse, human and zebrafish *ACRC* are expressed in adult somatic tissues (Uhlén et al. 2015). Considering the substantial mRNA expression levels of *ACRC* in adult tissues of zebrafish and humans, we suggest it also plays a role beyond DPC repair in meiosis in the germline.

Comparison of the expression levels of *acrc* and *sprtn* during zebrafish embryonic development showed that both genes are highly expressed up to 6 hpf, but also that *acrc* is expressed at least three-fold more strongly than *sprtn* between 1 hpf and 48 hpf, suggesting that Acrc is the dominant DPCR protease at these stages (Figure 2C). This is supported by our results showing that Acrc is absolutely required during embryonic development (Figure 3).

We showed that the proteolytic function of Acrc is essential for embryonic survival, as the mutant line with the deletion of the catalytic glutamate (E451) and three downstream amino acids (*rbi5* allele, Figure 3) displayed embryonic lethality after 6 hpf and before 10 hpf (Figure 3C). Previously, it was observed that the absence of the entire Acrc protein is lethal in zebrafish (Bhargava et al. 2020), but it was not known which domains of the Acrc are responsible for this phenotype. Considering that *acrc* mutants fully recover after injection of *Acrc*-WT mRNA into embryos at the one-cell stage and reach adulthood without obvious defects, we suggest that Acrc is the only DPCR factor that can repair specific DPC lesions at those early developmental stages (Figure 7). Other DPC proteases or nucleolytic DPC repair pathways cannot compensate for the loss of Acrc activity at this stage. Thus, we conclude that the function of Acrc in the soma during development is essential for embryo survival. In contrast, Acrc is not essential in adults, as rescued *acrc* mutants develop normally (the rescue mRNA injected is only functional for a few days after injection).

**Figure 7.**
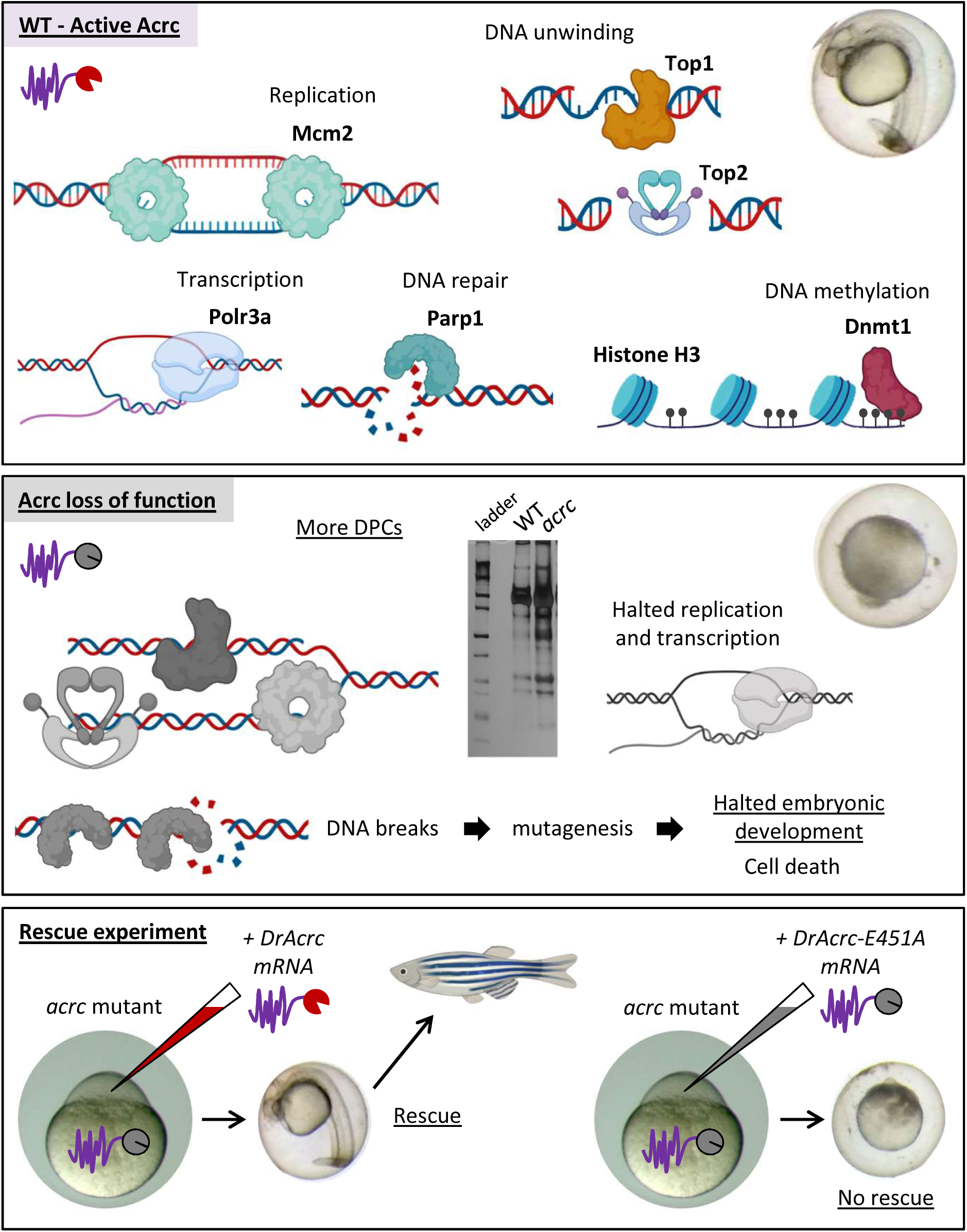
Model of proteolytic repair of DPCs by Acrc during embryonic development. Created with BioRender.com.

Our flexible non-genetic rescue system allows us to rapidly test different constructs encoding mutant forms or deletions of Acrc in a vertebrate model organism. Here, we used this system for the first time to examine the function of individual domains of Acrc. First, we confirmed the results from the *rbi5* mutant line showing that the catalytic function of Acrc is essential, because injection of *acrc* mRNA with the E451A mutation did not rescue the lethality phenotype (Figures 4 and S6). Previously, a similar experiment was performed in *Drosophila* using transgenic expression of WT and HE>AA mutant Acrc/Gcna in the mutant background (Bhargava et al. 2020) which showed that dsDNA breaks were not rescued by the HE>AA mutant, while egg hatching impairment was rescued, thereby indicating a separate function for Acrc/Gcna besides its catalytic activity. Using this separation-of-function approach, we found that deletion of the SprT-like domain impairs the function of Acrc and that the IDR domain alone cannot rescue the lethality phenotype (Figure 4), confirming that the proteolytic function of Acrc and not the IDR domain is essential for development. We additionally confirmed these findings by injecting mouse *Acrc* mRNA which consists only of an IDR region (Figure 4). Interestingly, an *Acrc* conditional KO mouse model showed defects in spermatogenesis including dsDNA breaks and defective chromatin compaction, possibly via the TOP2 interaction, but a direct role of mouse ACRC in DPCR has not yet been identified. Instead, it has been proposed that mouse ACRC promotes genome integrity during meiosis by mediating protein-protein interactions via its IDR domain and its SUMO-interaction motifs (Dokshin et al. 2020) and by acting as a histone chaperone (Ribeiro and Crossan 2022).

We showed that Sprtn and Acrc are not redundant, considering that *sprtn* is endogenously highly expressed during the onset of lethality in *acrc* mutants and the fact that injection of additional *sprtn* mRNA (Figure 2C) could not compensate for the lethality phenotype (Figure 4). Our results offer a direct functional proof that these two DPC proteases work in parallel pathways and that Sprtn cannot compensate for the loss of Acrc proteolytic activity. Because a catalytic mutation in the protease core causes early embryonic lethality, we investigated whether DPC accumulation precedes the lethality phenotype. Indeed, endogenous DPC levels were significantly increased in mutant embryos at 6 hpf before the onset of lethality (Figures 5, 6 and S7), suggesting a link between DPCR and the phenotype. Specifically, Top1-, Top2- and Parp1-DPCs hugely accumulate, followed by high accumulation of histone H3-DPCs, and moderate accumulation of Dnmt1-, Polr3a -and Mcm2-DPCs (Figures 6 and S7B-C). All the latter proteins are involved in essential cellular processes: TOP1, TOP2 and MCM2 in DNA replication, PARP1 in DNA repair, Pol III in transcription and DNMT1 in maintaining DNA methylation patterns following DNA replication (Figure 7). Considering very high accumulation of these and other DPCs as a consequence of Acrc inactivity and the fact that other factors including Sprtn, proteasome and nucleases cannot compensate for the lack of Acrc in the embryos, we conclude that ACRC is an essential DPC protease in the early vertebrate development (Figure 7). Cellular processes change tremendously during embryonic development: in zebrafish, the fertilized egg initially develops using maternally deposited mRNAs and proteins until the Maternal-Zygotic Transition (MZT) occurs around 3 – 4 hpf, leading to zygotic genome activation (ZGA) (Akdogan-Ozdilek et al. 2020). Additionally, during this time, cells divide every 30 minutes, which means that many DNA transactions occur simultaneously, such as DNA unwinding, transcription, replication and heterochromatin formation, which in turn makes DNA highly susceptible to DPC formation. Since this is a crucial time during vertebrate embryonic development, fidelity of DNA repair is absolutely essential.

## Materials and Methods

### Phylogeny and syntenic analysis

Nucleotide and protein sequences were retrieved from the NCBI (http://www.ncbi.nlm.nih.gov/) database using Blastx algorithm. Sequences were aligned with MUSCLE algorithm (Edgar 2004) and phylogenetic tree was constructed using Maximum Likelihood method in PhyML 3.0.1 software (Guindon and Gascuel 2003). Conserved synteny analysis between zebrafish and human ACRC was made using Genomicus (http://www.genomicus.biologie.ens.fr/genomicus), a conserved synteny browser synchronized with genomes from the Ensembl database (Louis et al. 2013).

### Model prediction

Domains and motifs were identified using SAPS and Motif Scan workspace (Brendel et al. 1992; Paysan-Lafosse et al. 2023; Sigrist et al. 2013). Protein topology schematics was visualized using IBS software (Liu et al. 2015). Models were build using Alphafold (Jumper et al. 2021; Varadi et al. 2022) and analyzed and visualized with UCSF Chimera (Pettersen et al. 2004). Disorder probability was predicted using PONDR-FIT (Xue et al. 2010).

### Zebrafish lines and handling

Adult wild-type (WT) zebrafish of the AB strain are kept at 28.5°C on a 14-hour light and 10-hour dark cycle under standard conditions (Aleström et al. 2020). Originally, adult zebrafish of both sexes of the AB strain were obtained from the European Zebrafish Resource Center at the Karlsruhe Institute of Technology (Germany). Zebrafish embryos were collected and kept in E3 medium (5mM NaCl; 0.17mM KCl; 0.33mM MgSO4; 0.33mM CaCl2) in Petri dishes at 28.5°C until 3 dpf under standard conditions. All handling and experiments were performed in accordance with the directions given in the EU Guide for the Care and Use of Laboratory Animals, Council Directive (86/609/EEC), and the Croatian Federal Act on the Protection of Animals (NN 135/06 and 37/13) under the project license HR-POK-023.

### Microinjections

Microinjections were performed using a microinjection system (Laboratory microinjector – FemtoJet® 4x series – Eppendorf) and needles (Eppendorf™ Femtotips™ Microinjection Capillary Tips). Microinjections in 1- to 4-cell stage embryos were performed by injecting 1 nl of premixed solutions into the yolk for the following purposes: 1) creating zebrafish *acrc* mutant lines using the CRISPR/Cas9 system, 2) rescue of the *acrc* mutant phenotype by mRNA injection. Embryos were then maintained in E3 medium under standard conditions at 28.5°C and staged as previously described (Kimmel et al. 1995).

### Creation of mutant zebrafish strains using CRISPR/Cas9 system, genotyping

Genetically modified zebrafish lines were created using Cas9 protein (EnGen® Spy Cas9 NLS, #M0646T, NEB) and previously established protocols combining PCR and reverse transcription to generate guide RNAs (Modzelewski et al. 2018). To create specific zebrafish *acrc* mutant strains, the following two guides targeting *acrc* exon 12 were designed using the CRISPRscan software (Moreno-Mateos et al. 2015):

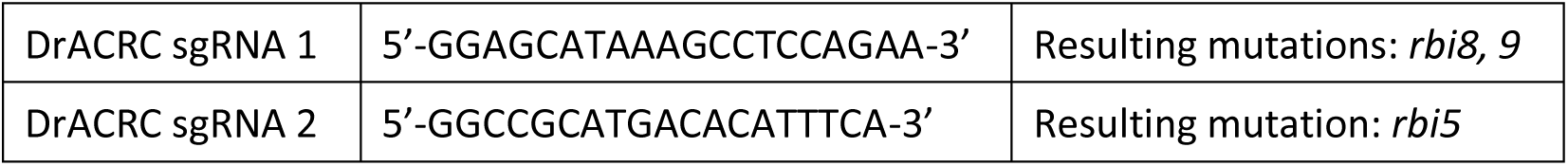

To introduce *acrc* mutations in exon 12, 1-cell stage zebrafish embryos were microinjected with 1 nl of the following solution: guide RNA (180 ng/µL), Cas9 protein (600 ng/µL), and KCl (300 mM). F0 injected embryos were raised and founders were later identified by sequencing of F1 progeny. For genotyping experiments, injected embryos were lysed with 0.5 mg/ml proteinase K (#J63710, AlphaAesar) in digestion buffer (10 mM Tris-HCl pH 8.5, 50 mM KCl, 0.3% Tween-20) for 3 hours at 55°C. PCR and sequencing was then performed using the following primer pair:

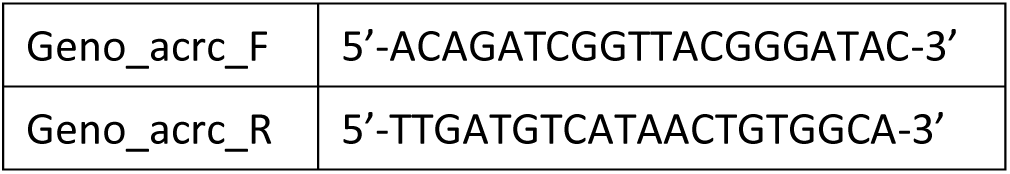

Three alleles were selected for further maintenance. All experiments were conducted with the *rbi5* allele which lacks 4 amino acids in the catalytic site of the SprT-like domain of Acrc (ΔEMCH 451-454). The mutant embryos with the other alleles (*rbi8* and *rbi9*, with 1 nt indel and 11 nt indel creating a premature stop at aa 477 and aa 472, respectively, see Figures 3 and S5A) displayed the same phenotype as *rbi5*.

### Rescue experiments: cloning and mRNA *in vitro* transcription

To perform rescue experiments, the WT coding sequence of zebrafish *gcna/acrc* (ENSDART00000169970.2) was amplified by PCR and cloned into the multicloning site of the pCS2+HisMyc vector (Rohr et al. 2006) between the XhoI and XbaI restriction sites and verified by sequencing. The vector contains an SP6 promoter, a multicloning site, a polyA tail and a unique restriction site after the polyA tag (KpnI). RNA was synthesized using purified, linearized plasmid by performing an *in vitro* transcription with SP6 using the Hiscribe SP6 RNA kit (#E2070S, NEB) and a cap analog from the ARCA kit (#S1411, NEB) for improved mRNA stability. The resulting RNA was purified using the RNA cleanup kit (#T2040, NEB) and injected into 1-2 cell stage *gcna/acrc* maternal mutant embryos or into control WT embryos (1nl of the mRNA solution: 250ng/μl RNA in 0.3M KCl and 0.015% phenol red). To determine functionally relevant amino acids and domains of *gcna/acrc*, we cloned deletion constructs by inverse PCR based on the original pCS2+HisMyc-DrAcrc plasmid using the following primer pairs (see Table). RNAs were then *in vitro* synthesized and injected as described above for WT *acrc*. Similarly, we obtained rescue constructs based on the same pCS2+HisMyc vector backbone with full-length coding sequences inserted between the XhoI and XbaI restriction sites for mouse *ACRC* (NM_001382234) or zebrafish *sprtn* (ENSDART00000158057.2) from Genosphere.

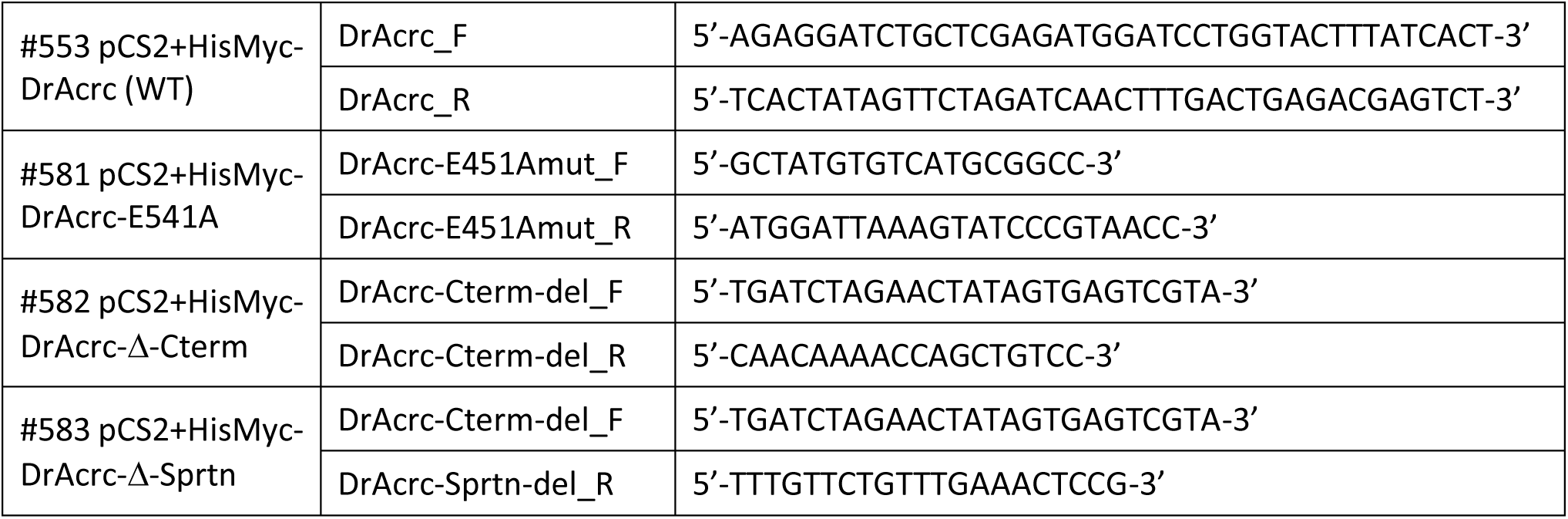

### Analysis of rescue experiments

To assess rescue efficiencies, the morphology of 1 dpf non-injected and injected embryos was assessed and embryos were categorized as follows: #1: “dead”; #2: “necrotic”: the yolk is intact, but the embryo is completely necrotic; #3: “very abnormal”: the embryo has a very short body axis due to necrosis and incomplete embryonic development; #4: “abnormal”: the embryo has a head and an elongated tail but can display defects such as cyclopia, which are typically the result of earlier necrosis; #5: “WT-like”: the embryo has a morphology indistinguishable from WT. Each rescue construct was injected in a total of at least 2 clutches of *acrc ^rbi5/rbi5^* and/or *acrc ^rbi8/9^*, and at least 50 embryos were categorized.

### RT-qPCR to verify the presence of the rescue constructs

To verify that the rescue RNAs are present in the injected embryos, pools of 5-20 embryos were collected at 1 or 2 dpf, total RNA was extracted using the Total RNA isolation kit (#T2010, NEB) following manufacturer’s instructions. Total RNA was then quantified using the Bio-Spec Nano Spectrophotometer and RNA integrity was determined by agarose gel electrophoresis. For each sample, purified total RNA was reverse transcribed using the ProtoScript II First Strand cDNA Synthesis Kit (#E6560, NEB) and a 1:1 mix of oligodT and random hexamer primers to obtain cDNA. Quantification of zebrafish gene expression was performed using the RT-qPCR method of relative quantification in technical triplicates on 5 ng cDNA per reaction using the following primers:

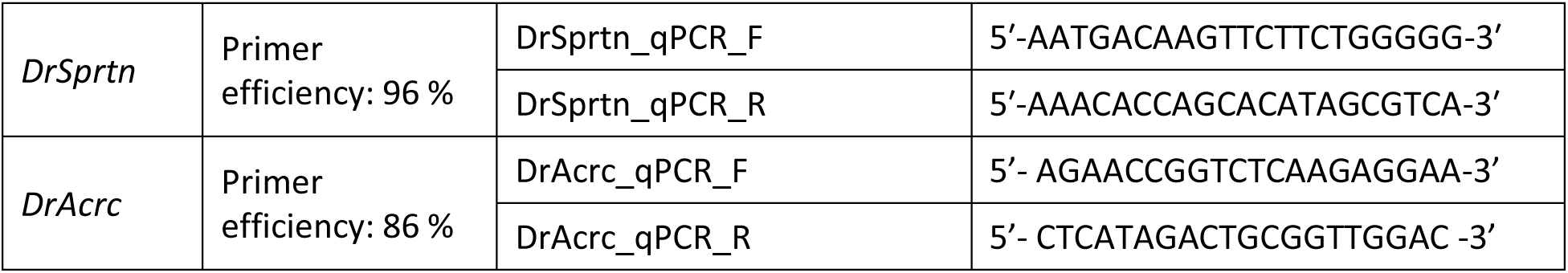
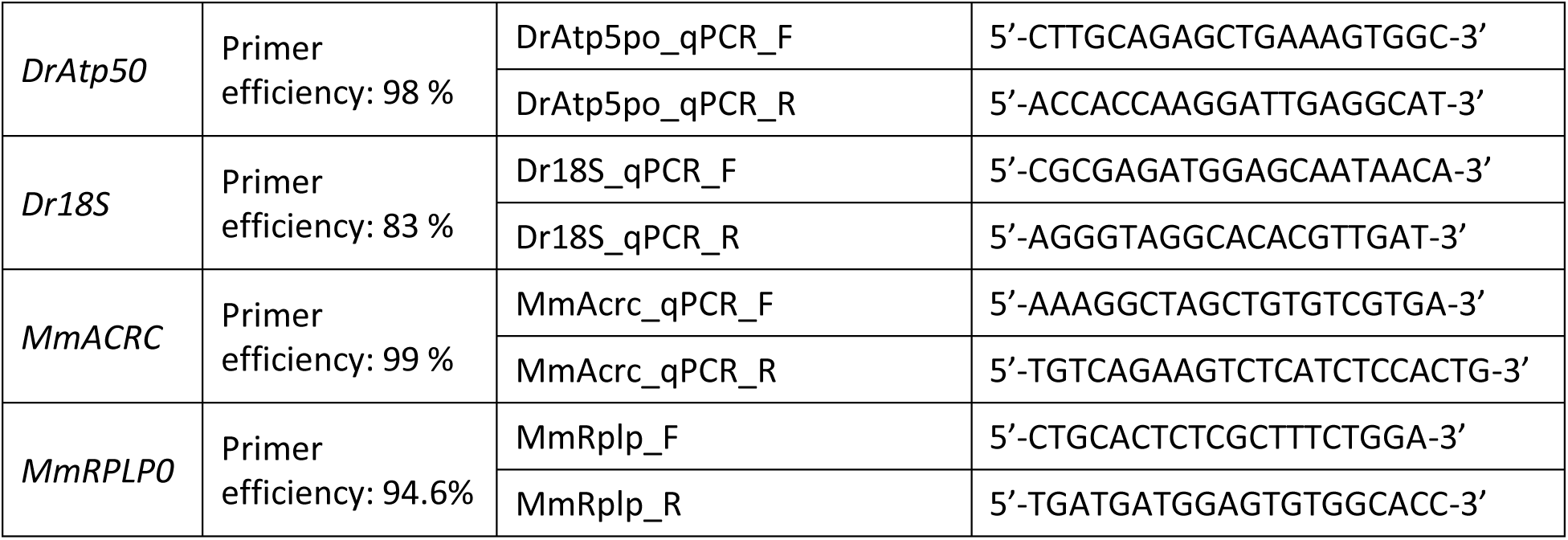

### Imaging

Zebrafish embryos were imaged using a Samsung 13-megapixel camera with an f/1.9 aperture applied to the ocular of a Motic SMZ-171 binocular.

### Zebrafish and mouse tissues collection

Adult male and female zebrafish were sacrificed for tissue collection according to regulations by immersion in ice-cold water supplied with 10% MS-222 (Tricaine) for 30 min and dissected to collect the following tissues: brain, liver, kidney, intestine, gonads (testes and ovaries), eye, gills, muscle. Tissues from n= 3 adults were pooled, put in RNALater and frozen. Wild-type zebrafish embryo samples were collected and dry-frozen at 1, 4, 6, 12, 24, 48, and 72 hpf: 3 pools of 30 embryos per condition. The following mouse tissues were donated by Prof. Balog (RBI, see acknowledgments): brain, liver, kidney, intestine, gonads (testes and ovaries), and processed similarly to the zebrafish tissues. The mouse tissues were derived from 3 females and 3 males, all of them 4 months old.

### RNA extraction and cDNA synthesis

Collected adult zebrafish and mouse tissue samples were thawed on ice and submitted to homogenization at 13500 min^-1^ for 30 seconds with Ultra turrax t25. Total RNA was then extracted from max. 50 µg tissue for each sample using the Total RNA isolation kit (#T2010, NEB) following manufacturer’s instructions. Total RNA was then quantified using the Bio-Spec Nano Spectrophotometer and RNA integrity was determined by agarose gel electrophoresis. For each sample, 1 µg of purified total RNA was reverse transcribed using the ProtoScript II First Strand cDNA Synthesis Kit (#E6560, NEB) and random hexamer primers to obtain 50 ng/µl cDNAs.

### Quantitative RT-PCRs

Quantification of zebrafish gene expression was performed using the RT-qPCR method of relative quantification in technical triplicates on 10 ng cDNA per reaction using the following primers:

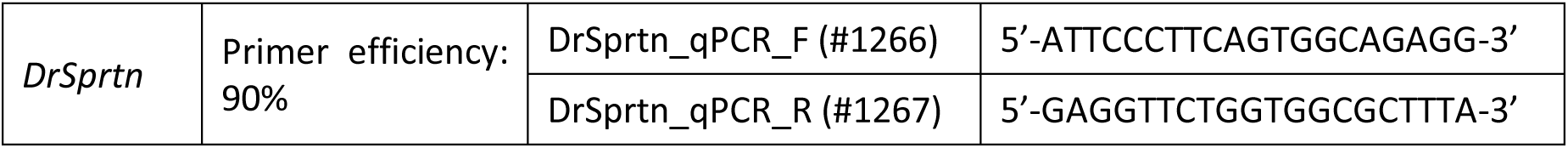
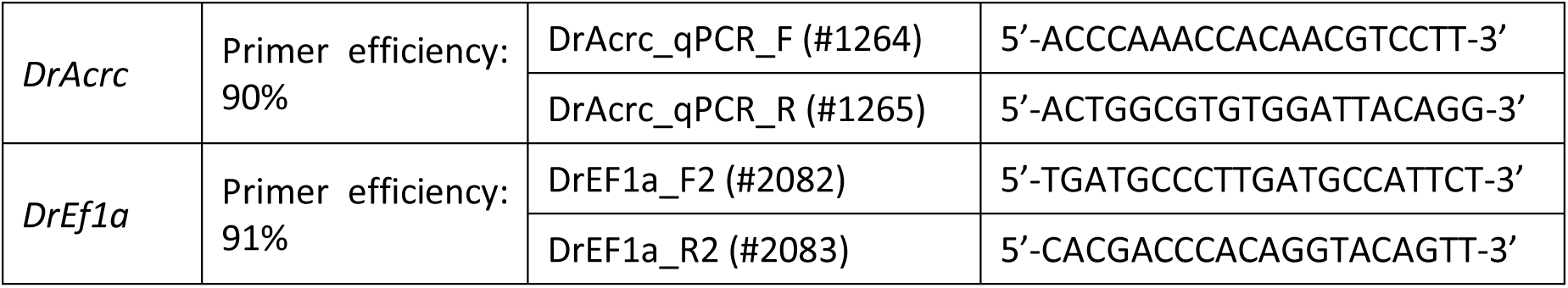

Quantification of mouse gene expression was performed using the RT-qPCR method of relative quantification in technical triplicates on 10 ng cDNA per reaction using the following primers:

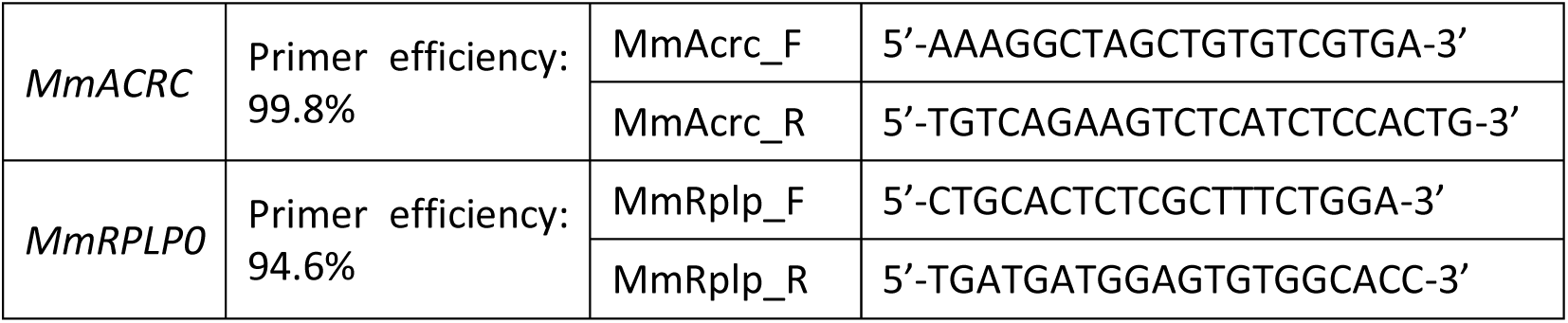

Expression of target genes was normalized to the housekeeping gene (HKG) *elongation factor 1α (eef1a1l1*) as previously described (Popovic et al. 2010). Mouse *Acrc* expression was normalized to the housekeeping gene Ribosomal Protein Lateral Stalk Subunit P0 (*Rplp0)*. The quantitative RT-PCRs were performed on a 7300 Real-Time PCR System (Applied Biosystems) using Power SYBR Green PCR Master Mix (#4367659, Applied Biosystems) according to manufacturer’s instructions. After initial denaturation at 95°C for 10 min, 40 amplification cycles were performed with denaturation at 95°C/15 s, annealing and elongation at 60°C/1 min, all together followed by melting curve analysis. Data analysis was performed using the 7500 Fast System SDS software (Applied Biosystems) followed by the relative quantification using Q-Gene method (Muller et al. 2002; Simon 2003) which is based on the following formula yielding the Mean Normalized Expression (MNE):

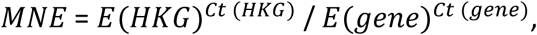

E (HKG) – amplification efficiency of the housekeeping gene, Ct (HKG) – average Ct value of the HKG for a particular tissue, E (gene) – amplification efficiency of the gene of interest, Ct (gene) – average Ct value of the gene of interest for that particular tissue. Data are given as MNE*10^6^ ± SD.

### Isolation and detection of DPCs by SDS-KCl method

Isolation of DPCs was performed using the SDS/KCl precipitation assay as previously described (Zhitkovich and Costa 1992; Mórocz et al. 2017). This method was used for DPC isolation from WT and *acrc^rbi8/9^* maternal mutant zebrafish embryos. Pools of 100 untreated embryos were collected at 6 hpf. Samples were then placed in liquid nitrogen and incubated overnight. The next day, samples were thawed in a thermoblock (55 °C/15 min) and then sonicated. Proteins were then precipitated by adding 400 μl KCl buffer (200 mM KCl; 20 mM Tris, pH 7.5) followed by incubation on ice for 5 min. The precipitated proteins were pelleted by centrifugation at 21,000 rcf for 5 min (4 °C), and the supernatant was kept for quantification of soluble DNA. The pellet was washed three times by adding 400 μl KCl buffer, followed by incubation at 55 °C for 5 min, 5 min on ice, and centrifugation at 15,000 rcf at 4 °C for 5 min. After washing, each pellet was resuspended in 400 μl KCl buffer containing proteinase K (0.2 mg/ml) and incubated at 55 °C for 1 h. After incubation, BSA (1.25 mg/ml) was added and samples were incubated on ice for 5 min. After centrifugation at 15,000 rcf at 4 °C for 5 min, the supernatant contained the cross-linked DNA. From both the soluble and cross-linked DNA, 50 μl of the sample was taken for treatment with RNase A (0.2 mg/ml) for 30 min at 37 °C. Soluble DNA and cross-linked DNA were quantified using the Quant-iT™ 1X dsDNA HS assay (ThermoFischer Scientific, MA, USA), and the amount of DPCs was calculated as the ratio of cross-linked DNA to total DNA (soluble + cross-linked).

### Isolation and detection of DPCs by RADAR method

DPC isolation and detection from zebrafish embryos was performed using the modified RADAR (rapid approach to DNA adduct recovery) assay which we previously adapted (Anticevic et al. 2023) from the original protocol developed by Kiianitsa and Maizels (2013). In brief, DPCs were isolated from 50 embryos (at 6 hpf) per sample using following steps: (1) lysis with pre-warmed lysis buffer (6 M guanidinium isothiocyanate, 10 mM Tris-HCl (pH 6.8), 20 mM EDTA, 4% Triton X-100, 1% N-lauroylsarcosine sodium and 5% β-mercaptoethanol) for 10 min at 55 ᵒC with vigorous mixing; (2) addition of an equal volume of 98% ethanol to precipitate DNA with crosslinked proteins; (3) centrifugation at 10,000 rcf for 10 min at 4 ᵒC; (4) washing the DNA pellet four times with wash buffer (20 mM Tris-HCl (pH 7.4), 1 mM EDTA, 50 mM NaCl, 50% EtOH) with the same centrifugation steps as in (3); (4) dissolving the pellet in 8 mM NaOH. A small aliquot of the DPC sample was set aside for DNA quantification and treated with proteinase K (20 mg/ml, Fisher Scientific, BP1700-100), and the remaining DNA was quantified using PicoGreen according to the manufacturer’s instructions (Invitrogen, P7581). After the DNA quantification of each sample, all DPC isolates were normalized to the same amount of DNA and treated with benzonase nuclease (Millipore, E1014) for 1 h at 37 °C, followed by snap-freezing in liquid nitrogen and subsequent overnight lyophilization: freeze-drying at -50 °C at 5 Pa vacuum using a FreeZone 2.5 lyophilizer (Labconco, USA). Lyophilized DPC samples were dissolved in SDS loading buffer (4 M urea, 62.5 mM Tris-HCl (pH 6.8), 1 mM EDTA, 2% SDS).

To detect total DPCs, dissolved DPC samples were resolved on SDS-PAGE gradient gel (5-18 %) and visualized by silver staining, according to the manufacturer’s protocol (Sigma Aldrich, PROTSIL1). Specific DPCs were detected by western blot (for histone H3) or by dot blot (for Dnmt1, Top2, Top1, Parp1, Polr3a and Mcm2) with protein-specific primary antibodies. Briefly, for western blot, DPCs were resolved on 5-18% SDS-PAGE gradient gels and transferred to PVDF membranes (Roche, 03010040001). Membranes were blocked with 5% low-fat milk in TBST buffer (10 mM Tris-HCl (pH 7.5), 15 mM NaCl, 0.02% Tween 20) for 2 h at room temperature and then incubated with primary antibodies in 2.5% BSA in TBST overnight at +4 °C, followed by an incubation with an HRP-labelled anti-rabbit secondary antibody (Sigma-Aldrich, #A0545, 1:100000) for 1 hour and washed three times for 10 min in TBST buffer. Detection was performed according to the manufacturer’s instructions for the ECL blotting substrate (Biorad, #1705061) and visualized using the ChemiDoc™ XRS+ System (Bio-Rad, #1708299). For the detection of histone H3, a DPC equivalent of 50 ng total DNA was subjected to western blot and immunostained with the anti-H3 antibody (Cell Signaling, #9715, 1:3000). Other specific DPCs (Dnmt1, Top2, Top1, Parp1, Polr3a and Mcm2) were detected by dot blot with protein-specific antibodies using the Bio-dot microfiltration device (Bio-Rad Laboratories, CA, USA). Briefly, for dot blot, 200 μL of DPC sample per dot was loaded onto the nitrocellulose membranes (GE Healthcare, 10-6000-02), and the samples were vacuumed using vacuum pump at 700 mbar. For the detection of Dnmt1, Top2, Top1, Parp1, Polr3a and Mcm2, a DPC equivalent of 25 ng total DNA was subjected to dot blot and immunostained with the following primary antibodies respectively: anti-DNMT1 (Cell Signaling, #5032, 1:1000), anti-TOP2A (Abcam, ab52934, 1:1000), anti-TOP1 (Santa Cruz, sc-271285, 1:1000), anti-PARP1 (Cell Signaling, #9532, 1:1000), anti- POLR3A (Cell Signaling, #12825, 1:1000) and anti-MCM2 (Cell Signaling, #3619, 1:1000). Other steps were performed as previously described for western blot. DNA detection was performed by applying 2 ng of DNA from each sample treated with proteinase K to a nylon membrane and immunostaining with the anti-dsDNA antibody (Abcam, #ab27156, 1:9000) using the dot plot apparatus as previously described. HRP-labelled anti-rabbit secondary antibody was used for anti-DNMT1, anti-TOP2A, anti-PARP1, anti-POLR3A and anti-MCM2 primary antibodies, while HRP-labelled anti-mouse secondary antibody (Sigma-Aldrich, #A9044, 1:100000) was used for anti-TOP1 and anti-dsDNA primary antibodies.

### Statistical Analysis

Data shown are means with SD or SEM. Data were statistically analyzed using the GraphPad Prism software (v.9.0.0, GraphPad) as described below. Changes were statistically significant with p-value < 0.05.

1. Figure 2A-B: quantification of relative gene expression levels for *acrc* and *sprtn* in adult zebrafish tissues normalized to *eef1a1l1* housekeeping gene expression: an unpaired Student’s t-test was performed for each tissue to determine whether gene expression is significantly dependent on gender. Differences were not significant, except for *acrc* levels in kidney (*: p-value = 0.0451) and in gonads (**: p-value = 0.0013).
2. Figure 2C: quantification of gene expression levels for *acrc* and *sprtn* during zebrafish embryonic development normalized to *eef1a1l1* housekeeping gene expression. Each gene was analyzed separately and One-way ANOVA analyses were followed by Sidak’s multiple comparisons tests. Gene expression levels were compared separately within the first 6 hours post fertilization (maternal mRNA contribution) and between 6 hpf – 72 hpf (zygotic gene expression). For *acrc*, expression levels significantly dropped between 1 hpf and 6 hpf (*: p-value = 0.0297), but no other changes were significant.
3. Figure 2D: quantification of relative gene expression levels for *Acrc* in adult mouse tissues normalized to *Rplp0* housekeeping gene expression: an unpaired Student’s t-test was performed for each tissue to determine whether gene expression is significantly dependent on gender. Differences were not significant, except in gonads (*: p-value = 0.0114).
4. Figure 5A: quantification of DPC levels (% of crosslinked DNA) as measured by the SDS- KCl method. In comparison to WT embryos, *acrc^rbi8/rbi9^* mutant embryos display a significant increase in total DPC levels at 6 hpf. An unpaired Student’s t-test was performed for 3 biological replicates, pools of 100 embryos each (*: p-value = 0.0254).
5. Figures 5C-D: quantification of total DPCs based on silver stainings shown here and in Figure S7A. In comparison to WT embryos, *acrc^rbi5/rbi5^* mutant embryos display a significant increase in total DPC levels at 6 hpf. An unpaired Student’s t-test was performed for 4 biological replicates, pools of 50 embryos each (****: p-value < 0.0001). The same stainings were subdivided according to molecular weight of DPCs (low: < 40 kDa, middle: 41-150 kDa, high: > 151 kDa) and quantified separately. An unpaired Student’s t-test was performed for 3 (HMW) or 4 biological replicates (MMW and LMW) (****: p-value < 0.0001).
6. Figures 6B and 6D: quantification of H3-DPC levels based on western blots shown here and in Figure S7C and quantification of Dnmt1-, Top2-, Top1-, Parp1-, Polr3a- and Mcm2- DPC levels based on dot blots shown here and in Figures S7B-C. In comparison to WT embryos, *acrc^rbi5/rbi5^* mutant embryos display a significant increase in Dnmt1-, Top2-, H3-, Top1-, Parp1-, Polr3a- and Mcm2-DPC levels at 6 hpf. An unpaired Student’s t-test was performed for 2 (Top2 and Polr3a) or 3 (Dnmt1, histone H3, Top1, Parp1 and Mcm2) biological replicates, pools of 50 embryos each (*: p-value < 0.05, ***: p-value < 0.001, ****: p-value < 0.0001).
7. Supplemental Figure S6B, Supplemental Table 3: quantifications of rescue levels: percentage of live embryos after 1 dpf: non-injected WT (n= 14 experiments, 681 embryos; 96.28 ± 5.97% alive), WT injected with DrAcrc-WT mRNA (n= 6 experiments, 289 embryos; 95.88 ± 2.41% alive), uninjected *acrc* maternal mutant embryos (n=9 experiments, 446 embryos; 1.52 ± 02.53% alive) and rescued *acrc* maternal mutant embryos with DrAcrc-WT mRNA (n=5 experiments, 495 embryos; 94.80 ± 3.41% alive). A 2-way ANOVA followed by Tukey’s test was performed. ****: p-value < 0.0001.

## Acknowledgments

Mouse tissues were a kind gift from Prof. Tihomir Balog, Ruder Boskovic Institute (RBI). The pCS2+HisMyc plasmid used for cloning the rescue constructs was a kind gift from Prof. Salim Seyfried, Potsdam University, Germany. The M.P. research group and this work was supported by the Croatian Science Foundation Installation Grant (UIP-2017-05-5258), the Slovenian- Croatian Bilateral Research Project grant (IPS-2020-01-4225), European Structural and Investment Funds STIM – REI project (KK.01.1.1.01.0003) and Croatian Science Foundation under the project number HRZZ-IP-2024-05-9425.

## Author contributions

CS-P, CO and MK created the mutant zebrafish lines. MP performed the phylogenetic, structural and domain analysis. CS-P performed syntenic analysis, and CS-P, CO and VM performed the mRNA expression experiments. CO performed the rescue experiments and phenotyping. IA optimized the RADAR protocol. MK performed DPC analyses. CO, MK and MP wrote the manuscript. MP designed and supervised the project.

## List of supplemental files

**Figure S1.**
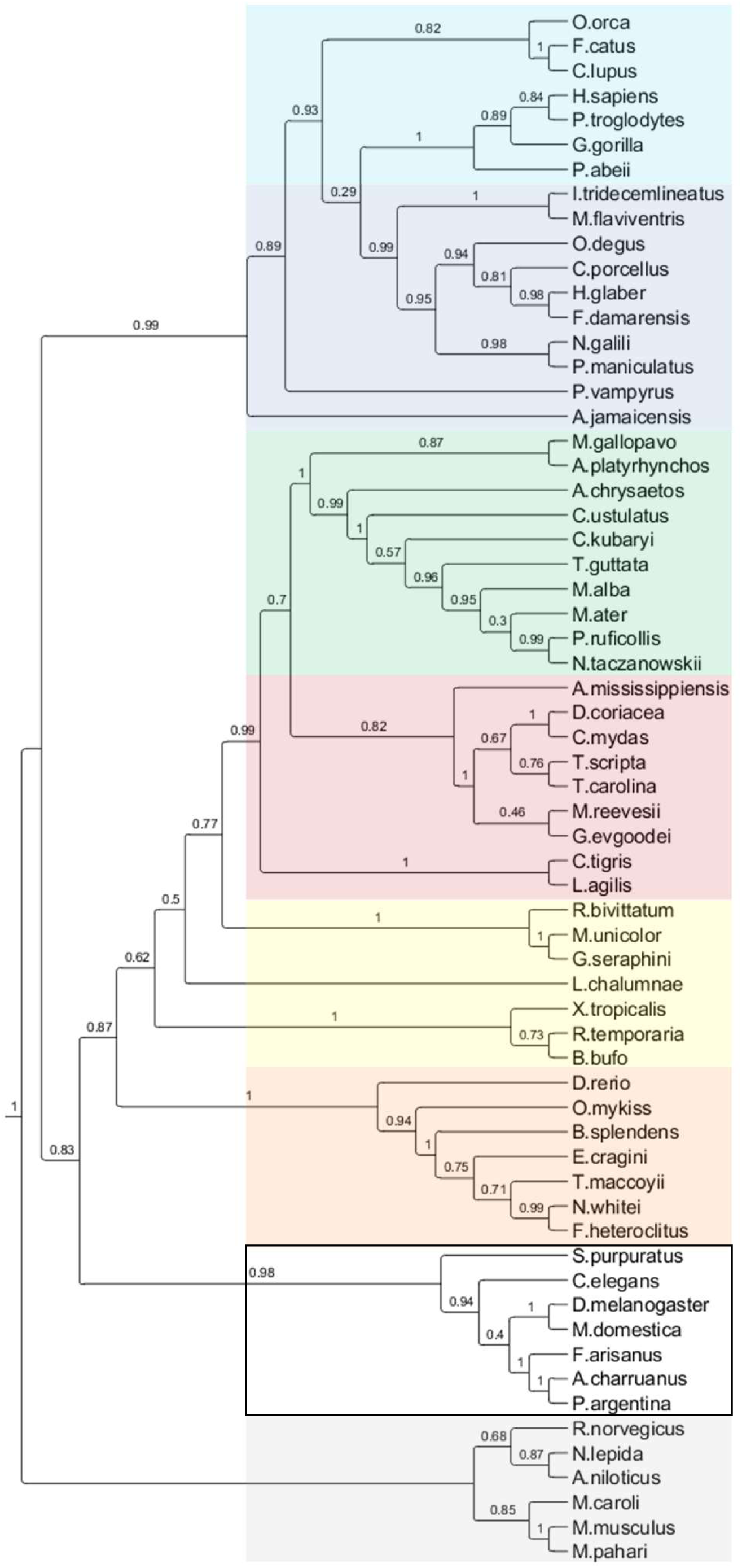
Extended phylogenetic tree of ACRC/GCNA orthologs based on the full-length protein sequences.

**Figure S2.**
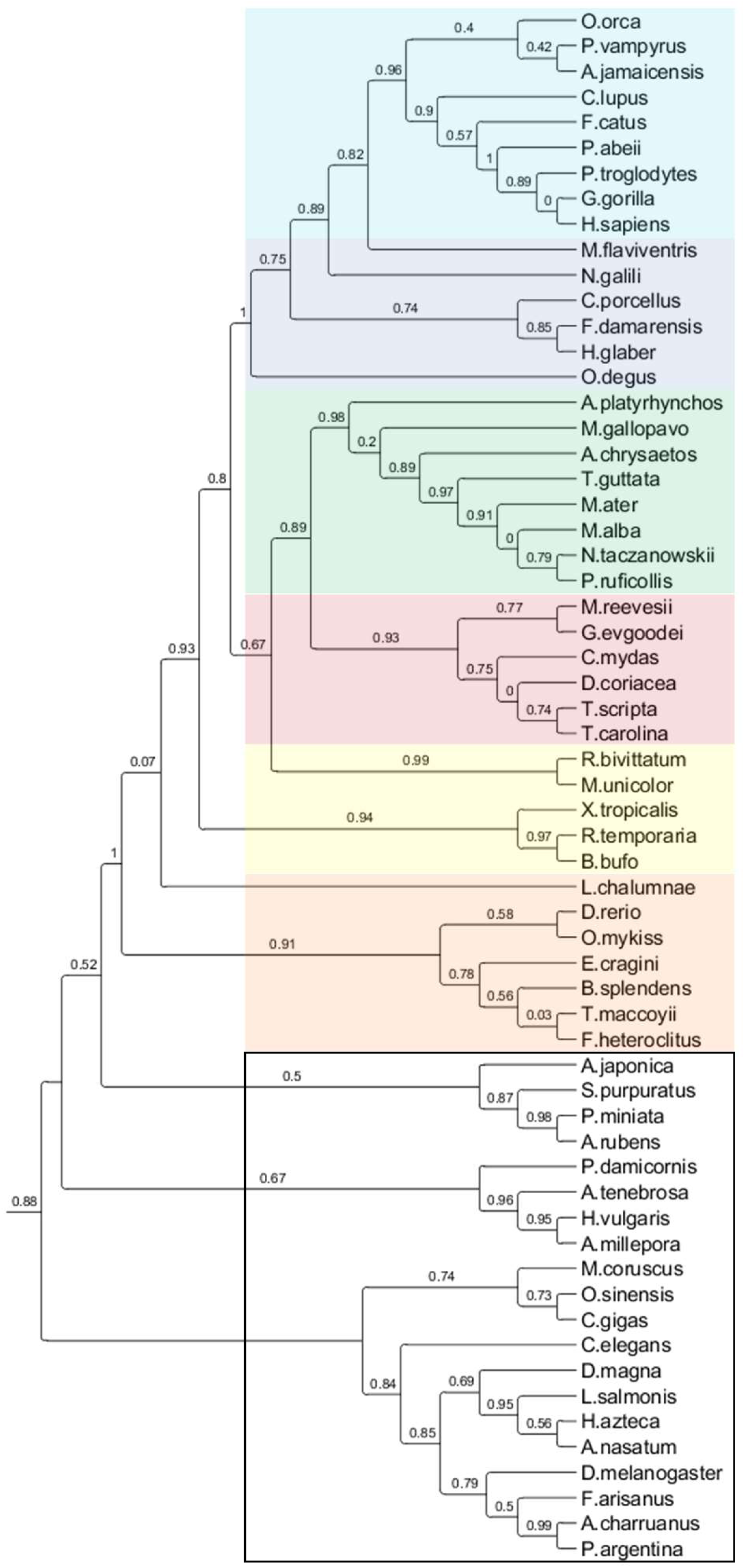
Extended phylogenetic tree of ACRC/GCNA orthologs based on the SprT-like domains.

**Figure S3.**
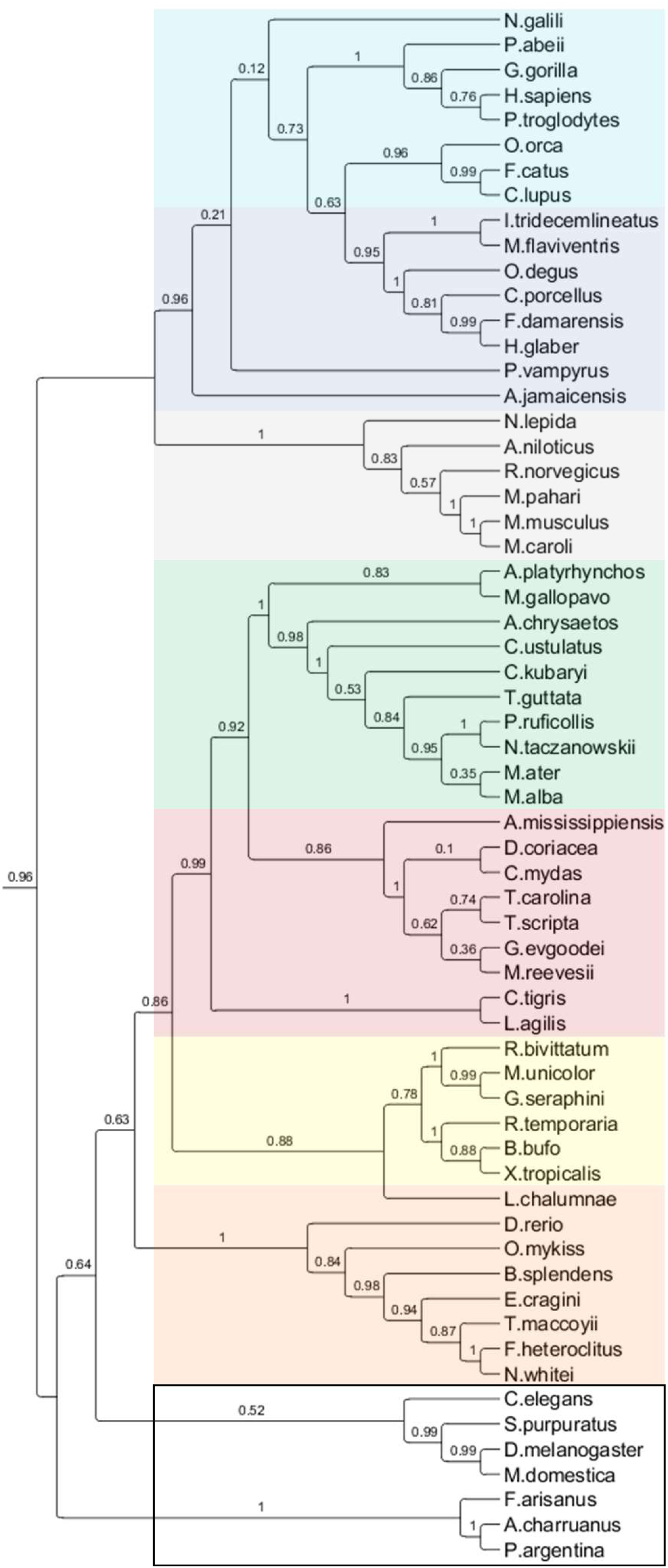
Extended phylogenetic tree of ACRC/GCNA orthologs based on the N-terminal / non-structured domains.

**Figure S4.**
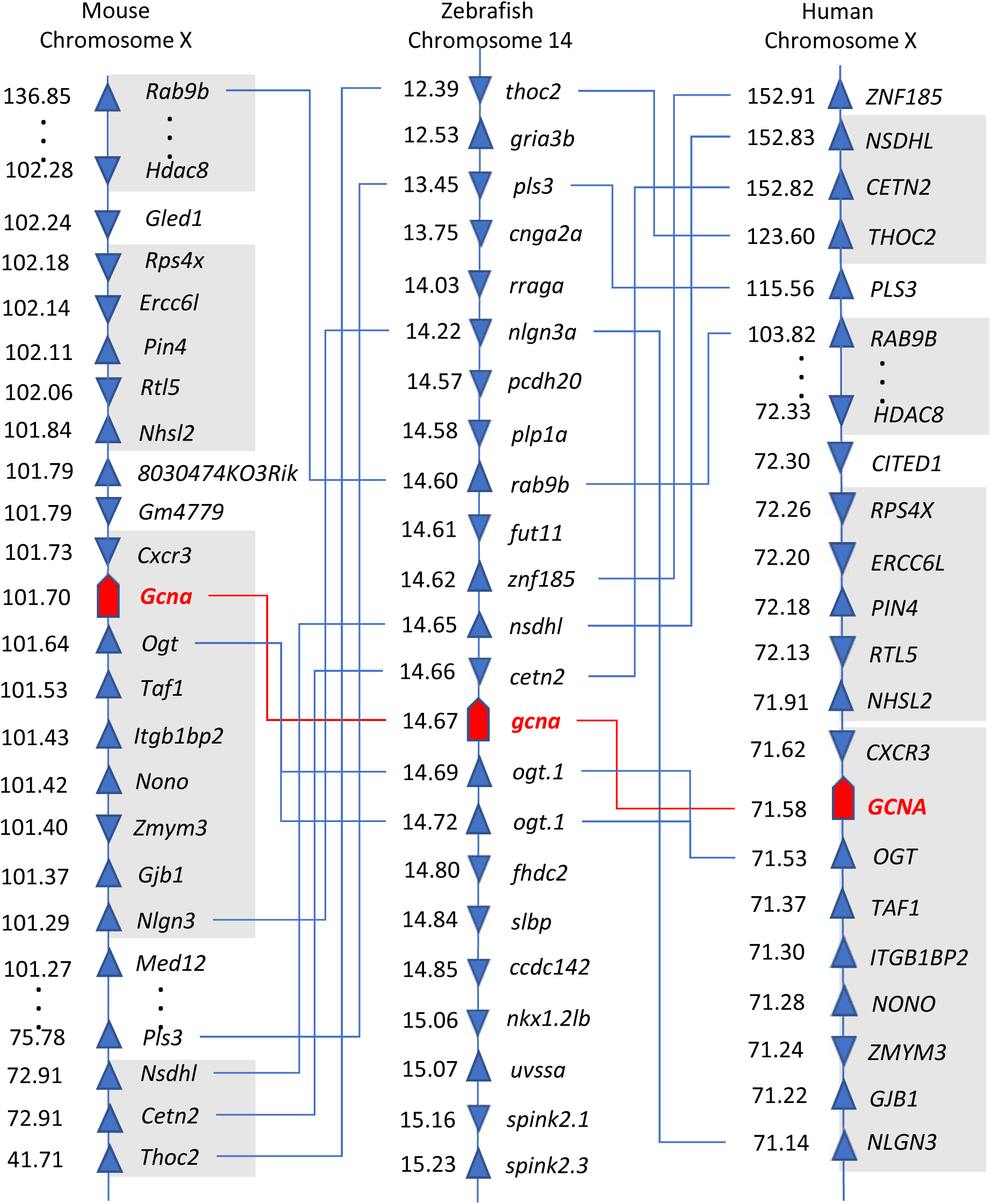
Conserved synteny analysis of zebrafish, mouse and human *ACRC/GCNA* genes.

**Figure S5.**
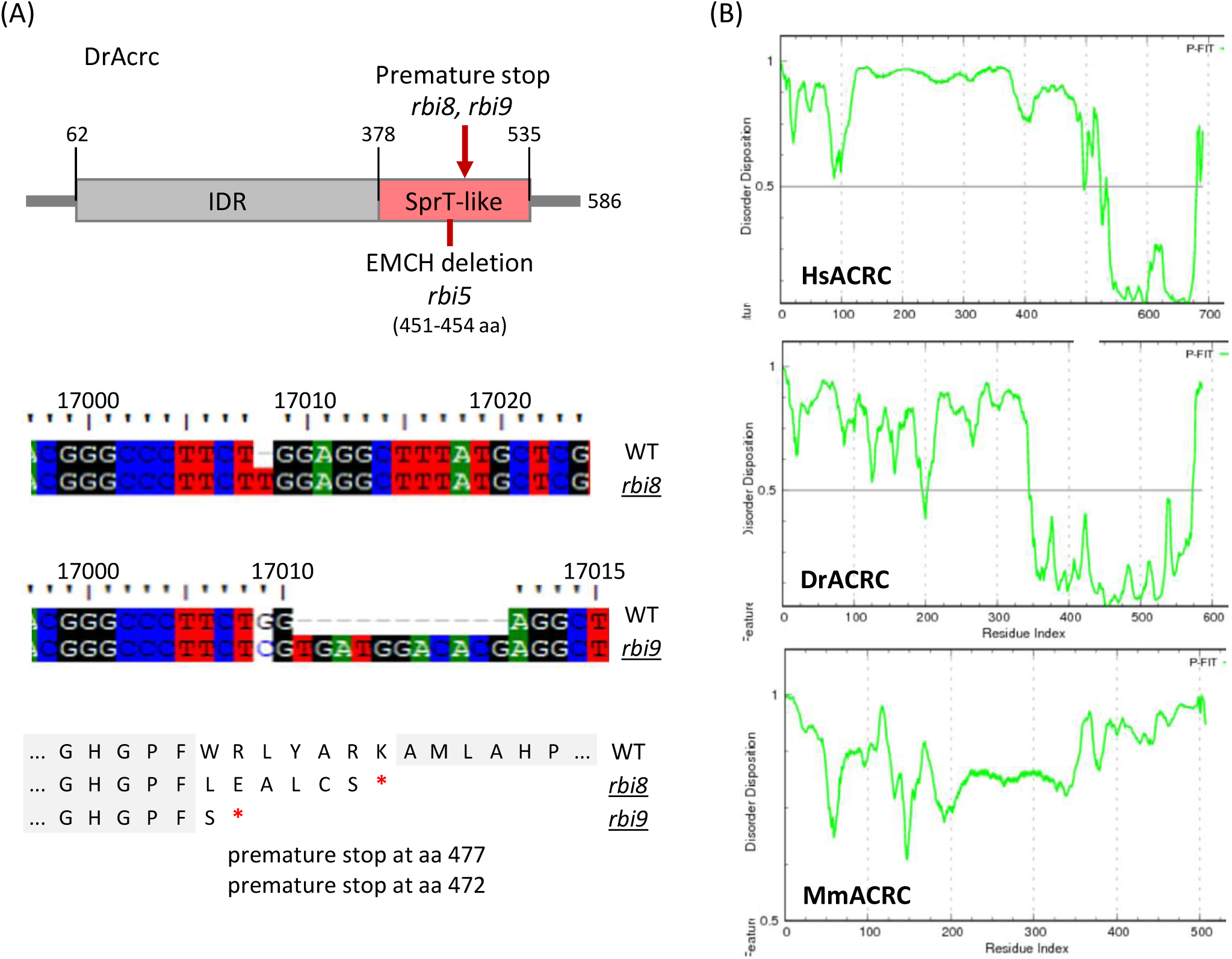
(A) Schematics showing the creation of two *acrc* mutant zebrafish strains; (B) Disorder disposition plot for human, zebrafish and mouse ACRC predicted using PONDR-FIT workspace.

**Figure S6.**
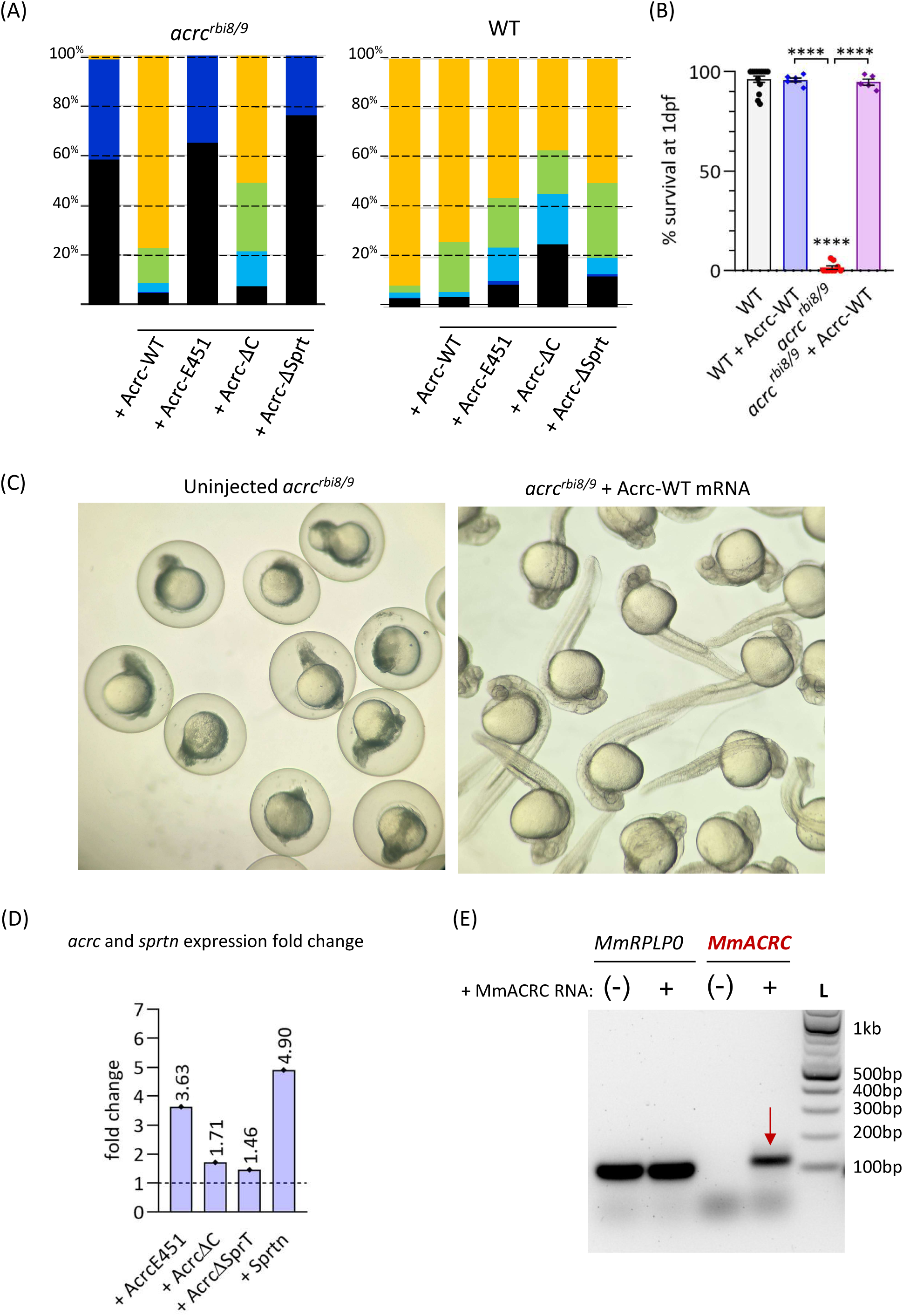
(A) Injection of Acrc-WT mRNA into *acrc^rbi8/9^* mutants fully rescues the lethal phenotype, while injection of Acrc-E451A mRNA does not (left panel). Ctrl: Injections of each constructs into the WT embryos are not toxic (right panel); (B) Quantification of A; (C) Images of uninjected *acrc^rbi8/9^* at 24 hpf (24 hours post fertilization) showing lethal phenotype and *acrc^rbi8/9^*embryos injected with Acrc-WT mRNA showing WT-like phenotype; (D) Expression levels of injected mRNA expressed as fold change to endogenous WT levels of each mRNA determined by RT-PCR, and ( E) mouse Acrc expression after injection determined by PCR (atp50 as a control, L-DNA ladder)

**Figure S7:**
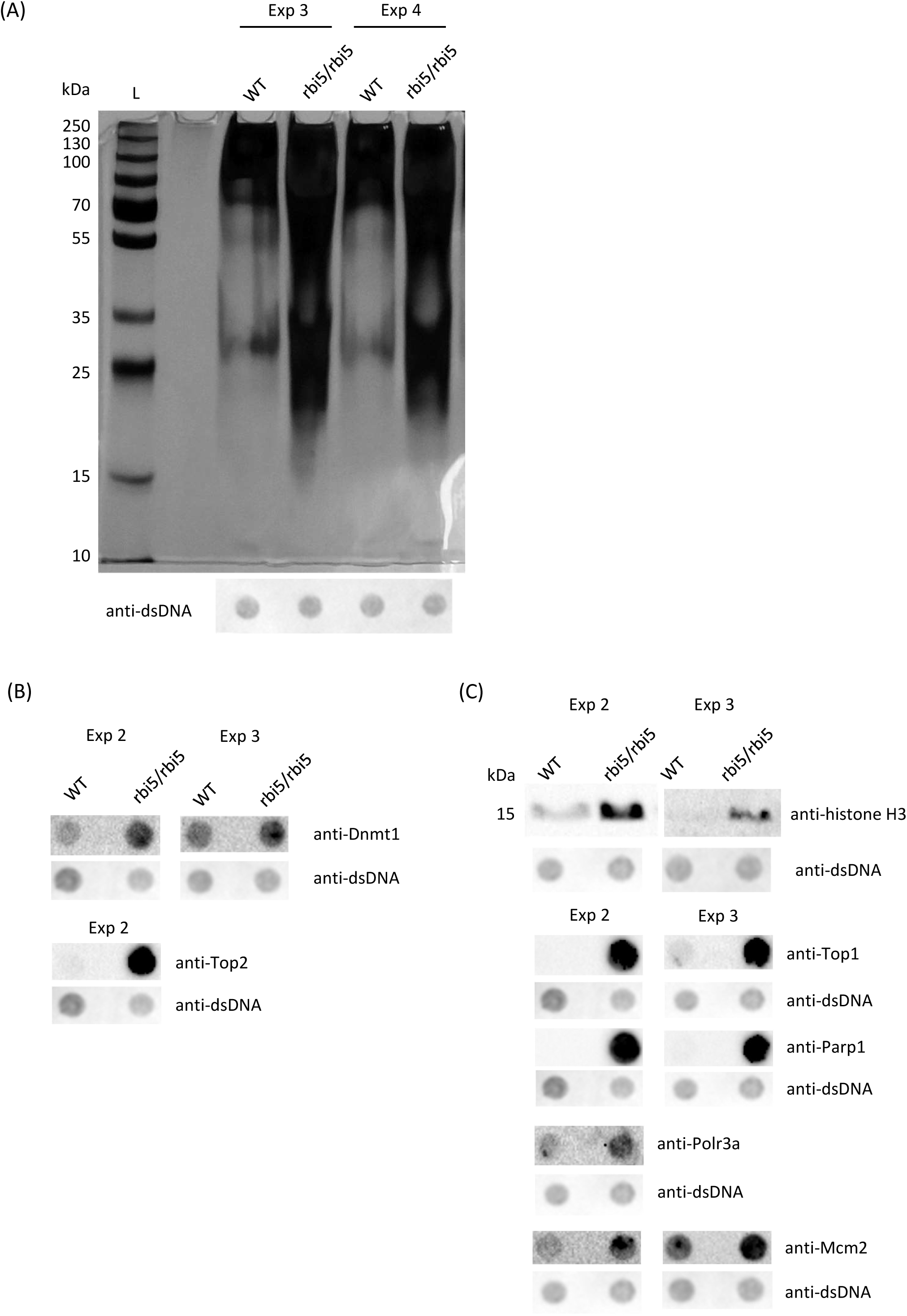
(A) Visualisation of DPCs in 6-hours old WT and *acrc^rbi5/rbi5^* mutant embryos isolated with RADAR, third and four biological replicates used for the quantification in Figures 5C-D; (B) Dnmt1-DPCs and Top2-DPCs detected with protein-specific antibodies using dot blot in the DPC isolates and (C) histone H3-, Top1-, Parp1-, Polr3a- and Mcm2-DPCs detected with protein-specific antibodies using dot blot in the DPC isolates. Second and third biological replicates used for the quantification in Figures 6B and 6D.

**Table S1.**
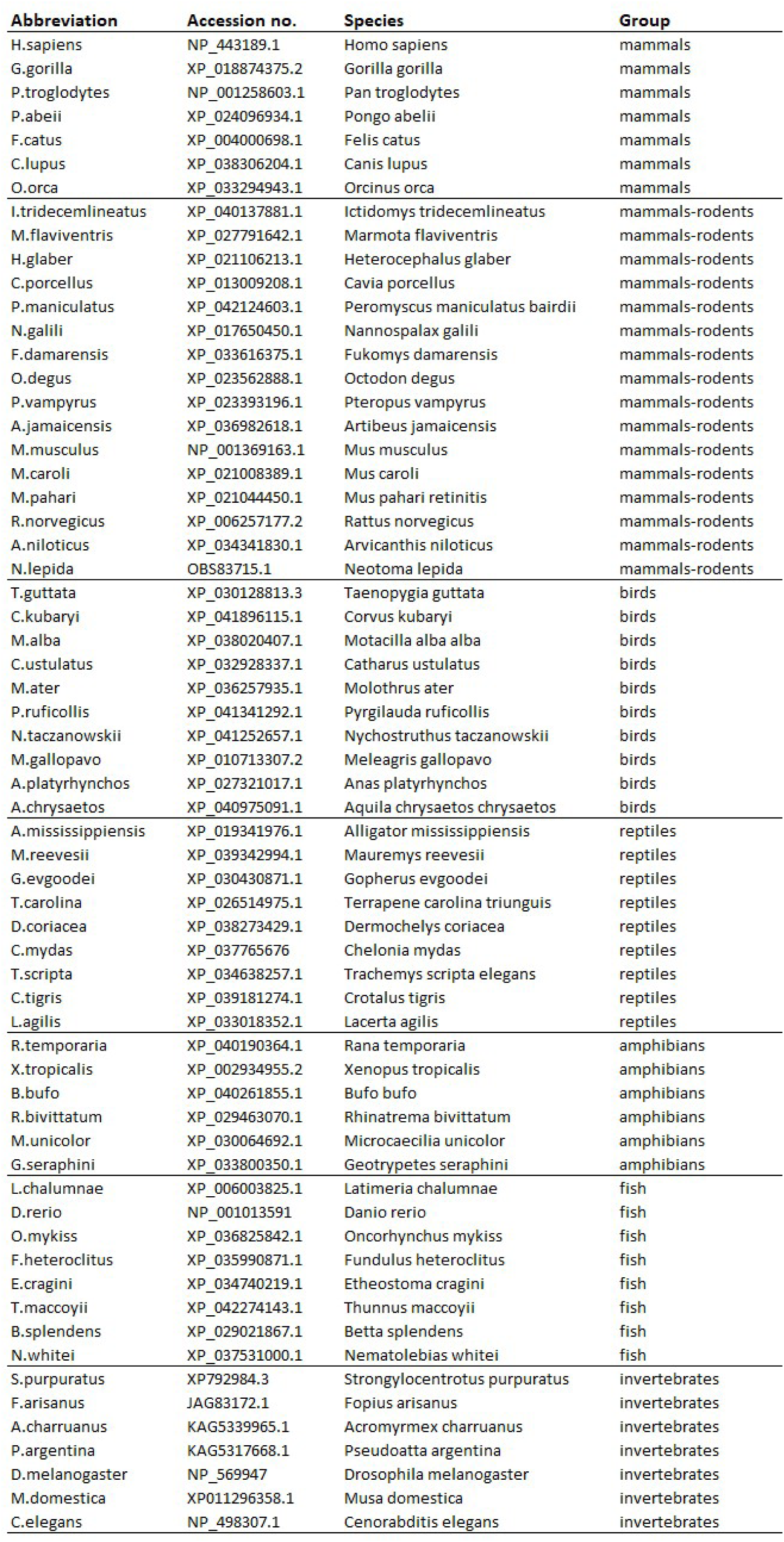
Protein sequences used for building the phylogenetic tree of full length ACRC/GCNA orthologs. Accession numbers in NCBI database, species abbreviations and group designations.

**Table S2.**
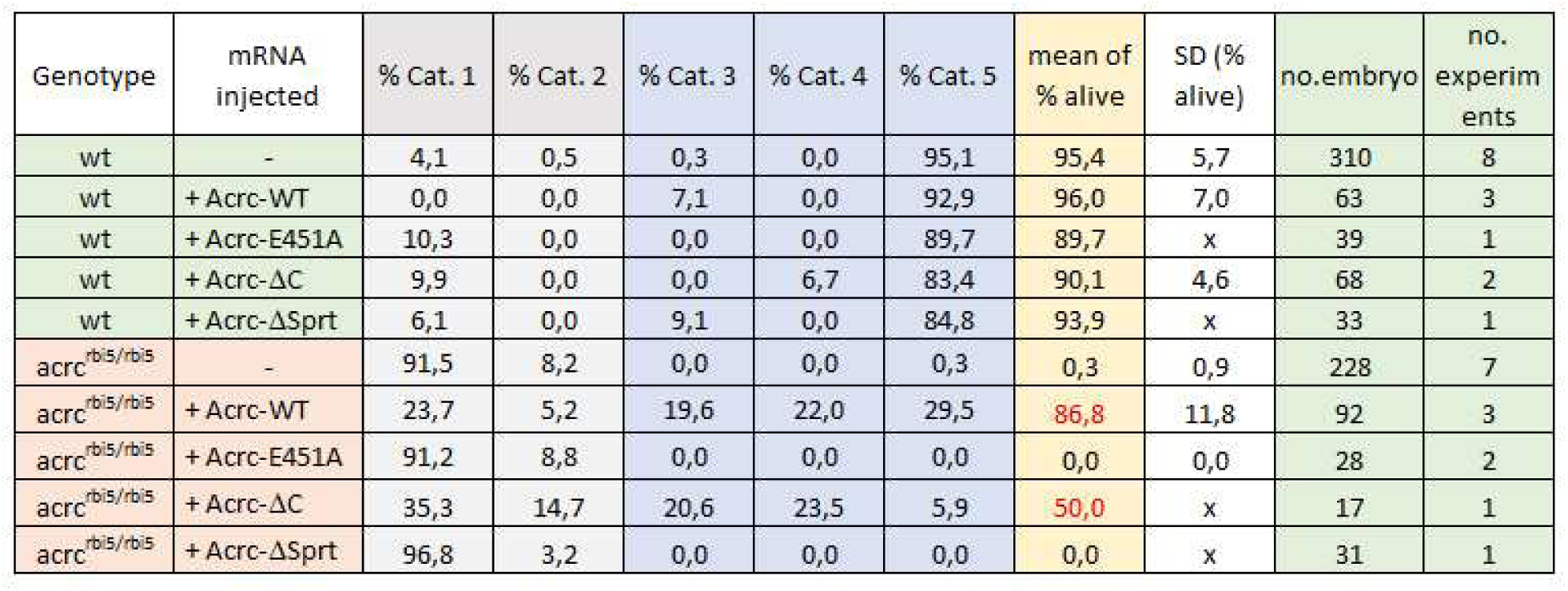
Rescue experiments with the DrAcrc rescue constructs Acrc-WT, Acrc-E451A, Acrc-ΔC and Acrc-ΔSprT injected in *acrc^rbi5/rbi5^* (ΔEMCH) maternal mutant fish. Raw data: numbers of embryos in each assessment category (Cat.) at 1 dpf. *acrc^rbi5/rbi5^*mutant line: percentage of embryos in each assessment category for each experiment.

**Table S3.**
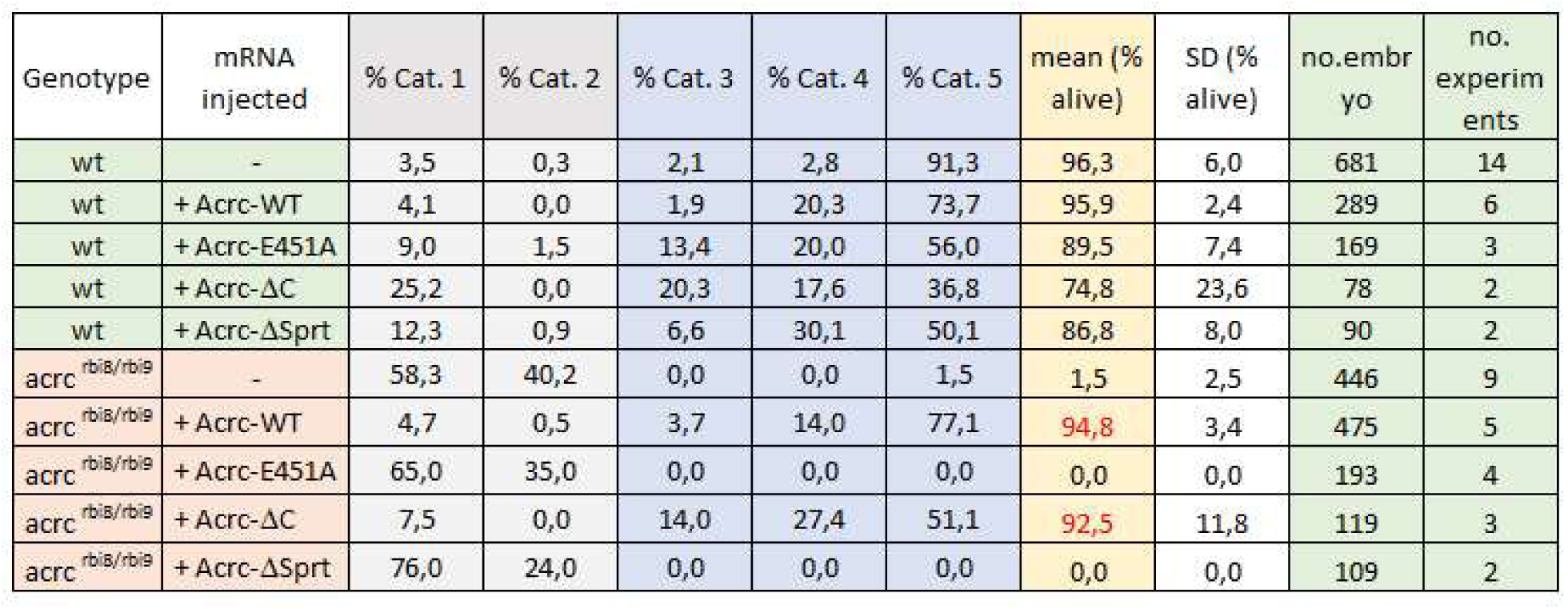
Rescue experiments with the DrAcrc rescue constructs Acrc-WT, Acrc-E451A, Acrc-ΔC and Acrc-ΔSprT in the *acrc^rbi8/9^* maternal mutant fish. Raw data: numbers of embryos in each assessment category at 1 dpf. Rbi8-9 mutant line: percentage of embryos in each assessment category for each experiment.

**Table S4.**
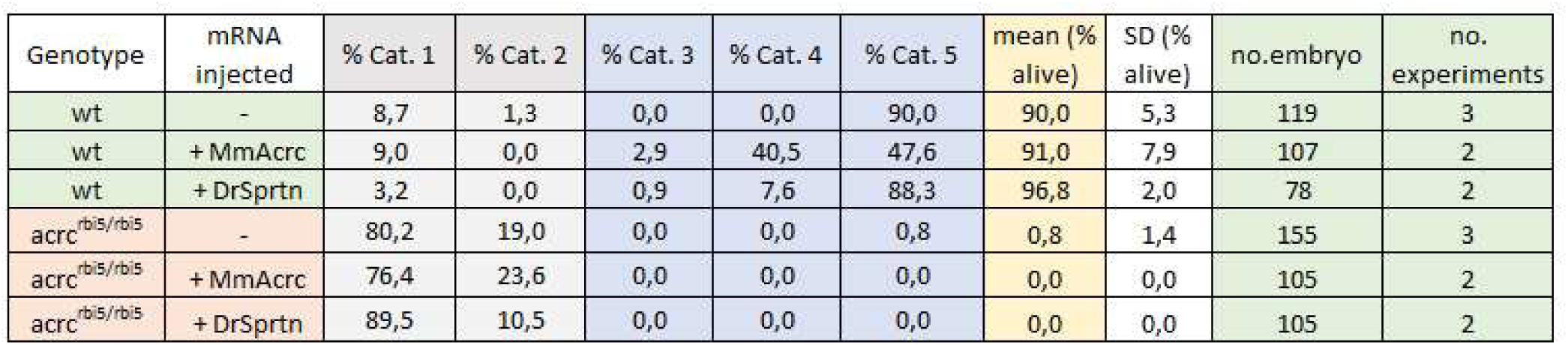
Rescue experiments with the DrSprtn and the MmAcrc rescue constructs in the *acrc* ΔEMCH maternal mutant fish. Raw data: numbers of embryos in each assessment category at 1 dpf. Rbi5 mutant line: percentage of embryos in each assessment category for each experiment.

