## Supplemental Table 1 for "ACRC/GCNA is an essential protease that repairs DNA-protein crosslinks during vertebrate development"

Table S1

| Abbreviation | Accession no. | Species | Group |
|---|---|---|---|
| H.sapiens | NP_443189.1 | Homo sapiens | mammals |
| G.gorilla | XP_018874375.2 | Gorilla gorilla | mammals |
| P.troglodytes | NP_001258603.1 | Pan troglodytes | mammals |
| P.abeii | XP_024096934.1 | Pongo abelii | mammals |
| F.catus | XP_004000698.1 | Felis catus | mammals |
| C.lupus | XP_038306204.1 | Canis lupus | mammals |
| O.orca | XP_033294943.1 | Orcinus orca | mammals |
| I.tridecemlineatus | XP_040137881.1 | Ictidomys tridecemlineatus | mammals-rodents |
| M.flaviventris | XP_027791642.1 | Marmota flaviventris | mammals-rodents |
| H.glaber | XP_021106213.1 | Heterocephalus glaber | mammals-rodents |
| C.porcellus | XP_013009208.1 | Cavia porcellus | mammals-rodents |
| P.maniculatus | XP_042124603.1 | Peromyscus maniculatus bairdii | mammals-rodents |
| N.galili | XP_017650450.1 | Nannospalax galili | mammals-rodents |
| F.damarensis | XP_033616375.1 | Fukomys damarensis | mammals-rodents |
| O.degus | XP_023562888.1 | Octodon degus | mammals-rodents |
| P.vampyrus | XP_023393196.1 | Pteropus vampyrus | mammals-rodents |
| A.jamaicensis | XP_036982618.1 | Artibeus jamaicensis | mammals-rodents |
| M.musculus | NP_001369163.1 | Mus musculus | mammals-rodents |
| M.caroli | XP_021008389.1 | Mus caroli | mammals-rodents |
| M.pahari | XP_021044450.1 | Mus pahari retinitis | mammals-rodents |
| R.norvegicus | XP_006257177.2 | Rattus norvegicus | mammals-rodents |
| A.niloticus | XP_034341830.1 | Arvicanthis niloticus | mammals-rodents |
| N.lepida | OBS83715.1 | Neotoma lepida | mammals-rodents |
| T.guttata | XP_030128813.3 | Taenopygia guttata | birds |
| C.kubaryi | XP_041896115.1 | Corvus kubaryi | birds |
| M.alba | XP_038020407.1 | Motacilla alba alba | birds |
| C.ustulatus | XP_032928337.1 | Catharus ustulatus | birds |
| M.ater | XP_036257935.1 | Molothrus ater | birds |
| P.ruficollis | XP_041341292.1 | Pyrgilauda ruficollis | birds |
| N.taczanowskii | XP_041252657.1 | Nychostruthus taczanowskii | birds |
| M.gallopavo | XP_010713307.2 | Meleagris gallopavo | birds |
| A.platyrhynchos | XP_027321017.1 | Anas platyrhynchos | birds |
| A.chrysaetos | XP_040975091.1 | Aquila chrysaetos chrysaetos | birds |
| A.mississippiensis | XP_019341976.1 | Alligator mississippiensis | reptiles |
| M.reevesii | XP_039342994.1 | Mauremys reevesii | reptiles |
| G.evgoodei | XP_030430871.1 | Gopherus evgoodei | reptiles |
| T.carolina | XP_026514975.1 | Terrapene carolina triunguis | reptiles |
| D.coriacea | XP_038273429.1 | Dermochelys coriacea | reptiles |
| C.mydas | XP_037765676 | Chelonia mydas | reptiles |
| T.scripta | XP_034638257.1 | Trachemys scripta elegans | reptiles |
| C.tigris | XP_039181274.1 | Crotalus tigris | reptiles |
| L.agilis | XP_033018352.1 | Lacerta agilis | reptiles |
| R.temporaria | XP_040190364.1 | Rana temporaria | amphibians |
| X.tropicalis | XP_002934955.2 | Xenopus tropicalis | amphibians |
| B.bufo | XP_040261855.1 | Bufo bufo | amphibians |
| R.bivittatum | XP_029463070.1 | Rhinatrema bivittatum | amphibians |
| M.unicolor | XP_030064692.1 | Microcaecilia unicolor | amphibians |
| G.seraphini | XP_033800350.1 | Geotrypetes seraphini | amphibians |
| L.chalumnae | XP_006003825.1 | Latimeria chalumnae | fish |
| D.rerio | NP_001013591 | Danio rerio | fish |
| O.mykiss | XP_036825842.1 | Oncorhynchus mykiss | fish |
| F.heteroclitus | XP_035990871.1 | Fundulus heteroclitus | fish |
| E.cragini | XP_034740219.1 | Etheostoma cragini | fish |
| T.maccoyii | XP_042274143.1 | Thunnus maccoyii | fish |
| B.splendens | XP_029021867.1 | Betta splendens | fish |
| N.whitei | XP_037531000.1 | Nematolebias whitei | fish |
| S.purpuratus | XP792984.3 | Strongylocentrotus purpuratus | invertebrates |
| F.arisanus | JAG83172.1 | Fopius arisanus | invertebrates |
| A.charruanus | KAG5339965.1 | Acromyrmex charruanus | invertebrates |
| P.argentina | KAG5317668.1 | Pseudoatta argentina | invertebrates |
| D.melanogaster | NP_569947 | Drosophila melanogaster | invertebrates |
| M.domestica | XP011296358.1 | Musa domestica | invertebrates |
| C.elegans | NP_498307.1 | Cenorabditis elegans | invertebrates |
