## Supplemental Table 2 for "ACRC/GCNA is an essential protease that repairs DNA-protein crosslinks during vertebrate development"

Table S2

| Genotype | mRNA injected | % Cat. 1 | % Cat. 2 | % Cat. 3 | % Cat. 4 | % Cat. 5 | mean of % alive | SD (% alive) | no.embryo | no. experiments |
|---|---|---|---|---|---|---|---|---|---|---|
| wt | - | 4,1 | 0,5 | 0,3 | 0,0 | 95,1 | 95,4 | 5,7 | 310 | 8 |
| wt | + Acrc-WT | 0,0 | 0,0 | 7,1 | 0,0 | 92,9 | 96,0 | 7,0 | 63 | 3 |
| wt | + Acrc-E451A | 10,3 | 0,0 | 0,0 | 0,0 | 89,7 | 89,7 | x | 39 | 1 |
| wt | + Acrc-ΔC | 9,9 | 0,0 | 0,0 | 6,7 | 83,4 | 90,1 | 4,6 | 68 | 2 |
| wt | + Acrc-ΔSprt | 6,1 | 0,0 | 9,1 | 0,0 | 84,8 | 93,9 | x | 33 | 1 |
| $acrc^{rbi5/rbi5}$ | - | 91,5 | 8,2 | 0,0 | 0,0 | 0,3 | 0,3 | 0,9 | 228 | 7 |
| $acrc^{rbi5/rbi5}$ | + Acrc-WT | 23,7 | 5,2 | 19,6 | 22,0 | 29,5 | 86,8 | 11,8 | 92 | 3 |
| $acrc^{rbi5/rbi5}$ | + Acrc-E451A | 91,2 | 8,8 | 0,0 | 0,0 | 0,0 | 0,0 | 0,0 | 28 | 2 |
| $acrc^{rbi5/rbi5}$ | + Acrc-ΔC | 35,3 | 14,7 | 20,6 | 23,5 | 5,9 | 50,0 | x | 17 | 1 |
| $acrc^{rbi5/rbi5}$ | + Acrc-ΔSprt | 96,8 | 3,2 | 0,0 | 0,0 | 0,0 | 0,0 | x | 31 | 1 |
