## Supplemental Table 3 for "ACRC/GCNA is an essential protease that repairs DNA-protein crosslinks during vertebrate development"

Table S3

| Genotype | mRNA injected | % Cat. 1 | % Cat. 2 | % Cat. 3 | % Cat. 4 | % Cat. 5 | mean (% alive) | SD (% alive) | no.embryo | no. experiments |
|---|---|---|---|---|---|---|---|---|---|---|
| wt | - | 3,5 | 0,3 | 2,1 | 2,8 | 91,3 | 96,3 | 6,0 | 681 | 14 |
| wt | + Acrc-WT | 4,1 | 0,0 | 1,9 | 20,3 | 73,7 | 95,9 | 2,4 | 289 | 6 |
| wt | + Acrc-E451A | 9,0 | 1,5 | 13,4 | 20,0 | 56,0 | 89,5 | 7,4 | 169 | 3 |
| wt | + Acrc-ΔC | 25,2 | 0,0 | 20,3 | 17,6 | 36,8 | 74,8 | 23,6 | 78 | 2 |
| wt | + Acrc-ΔSprt | 12,3 | 0,9 | 6,6 | 30,1 | 50,1 | 86,8 | 8,0 | 90 | 2 |
| $acrc^{rbi8/rbi9}$ | - | 58,3 | 40,2 | 0,0 | 0,0 | 1,5 | 1,5 | 2,5 | 446 | 9 |
| $acrc^{rbi8/rbi9}$ | + Acrc-WT | 4,7 | 0,5 | 3,7 | 14,0 | 77,1 | 94,8 | 3,4 | 475 | 5 |
| $acrc^{rbi8/rbi9}$ | + Acrc-E451A | 65,0 | 35,0 | 0,0 | 0,0 | 0,0 | 0,0 | 0,0 | 193 | 4 |
| $acrc^{rbi8/rbi9}$ | + Acrc-ΔC | 7,5 | 0,0 | 14,0 | 27,4 | 51,1 | 92,5 | 11,8 | 119 | 3 |
| $acrc^{rbi8/rbi9}$ | + Acrc-ΔSprt | 76,0 | 24,0 | 0,0 | 0,0 | 0,0 | 0,0 | 0,0 | 109 | 2 |
