## Supplemental Table 4 for "ACRC/GCNA is an essential protease that repairs DNA-protein crosslinks during vertebrate development"

Table S4

| Genotype | mRNA injected | % Cat. 1 | % Cat. 2 | % Cat. 3 | % Cat. 4 | % Cat. 5 | mean (% alive) | SD (% alive) | no.embryo | no. experiments |
|---|---|---|---|---|---|---|---|---|---|---|
| wt | - | 8,7 | 1,3 | 0,0 | 0,0 | 90,0 | 90,0 | 5,3 | 119 | 3 |
| wt | + MmAcrc | 9,0 | 0,0 | 2,9 | 40,5 | 47,6 | 91,0 | 7,9 | 107 | 2 |
| wt | + DrSprtn | 3,2 | 0,0 | 0,9 | 7,6 | 88,3 | 96,8 | 2,0 | 78 | 2 |
| $acrc^{rbi5/rbi5}$ | - | 80,2 | 19,0 | 0,0 | 0,0 | 0,8 | 0,8 | 1,4 | 155 | 3 |
| $acrc^{rbi5/rbi5}$ | + MmAcrc | 76,4 | 23,6 | 0,0 | 0,0 | 0,0 | 0,0 | 0,0 | 105 | 2 |
| $acrc^{rbi5/rbi5}$ | + DrSprtn | 89,5 | 10,5 | 0,0 | 0,0 | 0,0 | 0,0 | 0,0 | 105 | 2 |
